# Longitudinal cell division is associated with single mutations in the FtsZ-recruiting SsgB in *Streptomyces*

**DOI:** 10.1101/860916

**Authors:** Xiansha Xiao, Joost Willemse, Patrick Voskamp, Xinmeng Li, Meindert Lamers, Jan Pieter Abrahams, Navraj Pannu, Gilles P. van Wezel

**Author notes:** Corresponding authors. J.P. Abrahams; G.P. van Wezel. these authors contributed equally to the work (Xiansha Xiao, Joost Willemse).

## Abstract

In most bacteria, cell division begins with the polymerization of the GTPase FtsZ at the mid-cell, which recruits the division machinery to initiate cell constriction. In the filamentous bacterium *Streptomyces*, cell division is positively controlled by SsgB, which recruits FtsZ to the future septum sites and promotes Z-ring formation. Here we show via site-saturated mutagenesis that various amino acid substitutions in the highly conserved SsgB protein result in the production of ectopically placed septa, that sever spores diagonally or along the long axis, perpendicular to the division plane. Ectopic septa were especially prominent when cells expressed SsgB variants with substitutions in residue E120. Biochemical analysis of SsgB variant E120G revealed that its interaction with - and polymerization of - FtsZ had been maintained. The crystal structure of *S. coelicolor* SsgB was resolved and the position of residue E120 suggests its requirement for maintaining the proper angle of helix α3, thus providing a likely explanation for the aberrant septa formed in SsgB E120 substitution mutants. Taken together, our work presents the first example of longitudinal division in a free living bacterium, which is explained entirely by changes in the FtsZ-recruiting protein SsgB.

## INTRODUCTION

Bacterial cell division is mediated via the formation of a contractile ring that consists of the tubulin homologue FtsZ. FtsZ polymerizes into a ring-like structure (the Z-ring) which serves as a scaffold for the recruitment of other members of the cell division machinery or divisome (Adams and Errington, 2009). The GTP-dependent polymerization of FtsZ is essential for the constrictive force. Cryo-electron microscopy and cryo-electron tomography data suggest that the Z-ring consists of small discontinuous and single-layered filaments form a continuous ring through lateral association (Szwedziak *et al.*, 2014). By filament sliding and a hinge-opening conformational switch, the Z-ring drives constriction of the cell membrane, as shown by in vitro reconstitution and structural studies (Li *et al.*, 2013; Szwedziak *et al.*, 2014). Septum-site localization in unicellular bacteria depends on FtsA and ZipA that anchor FtsZ polymers to the cell membrane, with ZapA stabilizing the FtsZ filaments and promoting lateral interactions (Hale and deBoer, 1997; Pichoff and Lutkenhaus, 2002, 2005). In *Escherichia coli*, control of Z-ring timing and localization is governed by the Min system and by nucleoid occlusion, which negatively regulate FtsZ polymerization (Adams and Errington, 2009; Lutkenhaus, 2007; Margolin, 2005; Shapiro *et al.*, 2009).

Due to its central role in cell division, FtsZ is essential in nearly all bacteria. Two exceptions are the parasite *Mycoplasma* (Lluch-Senar et al., 2010), which has a reduced genome size and no cell walls, and *Streptomyces* (McCormick *et al.*, 1994; McCormick, 2009). Streptomycetes are filamentous Gram-positive bacteria in the phylum of Actinobacteria, that have a complex mycelial life style (Barka *et al.*, 2016). These bacteria produce over 60% of all known antibiotics and many other bioactive natural products (Hopwood, 2007; van der Heul *et al.*, 2018). Streptomycetes are model organisms for the study of multicellularity and bacterial morphogenesis (Claessen *et al.*, 2014; Flärdh and Buttner, 2009). Exponential growth of the multi-nucleoid vegetative hyphae is achieved by apical growth and branching. At this stage of the life cycle, cell division does not affect physical separation of the cells, but instead long syncytial cells are formed that are separated by cross-walls (Wildermuth and Hopwood, 1970). When the developmental programme is switched on, streptomycetes produce aerial hyphae, that ultimate differentiate into chains of unigenomic spores. Recent studies highlight that local membrane synthesis and branching may be an important divisome-independent mechanism for cell proliferation in *Streptomyces* (Celler *et al.*, 2016; Yagüe *et al.*, 2016).

Streptomycetes lack the canonical cell-division regulation systems such as Min and Noc (Jakimowicz and van Wezel, 2012). Instead, a positive control system has evolved, whereby FtsZ is actively recruited to the septum sites by SsgB, in concert with its paralogue SsgA (Willemse *et al.*, 2011). SsgA and SsgB belong to the SsgA-like proteins (SALPs), a family of regulatory proteins that is unique in sporulating actinobacteria, of which SsgA and SsgB are required for sporulation-specific cell division in *Streptomyces* (Noens *et al.*, 2005; Traag and van Wezel, 2008). Actinobacteria that form single spores, such as *Micromonospora* or *Thermobifida*, only have one SALP, namely the FtsZ-recruiting protein SsgB, while up to 14 SALPs can be found in those genera which form chains of spores, such as *Streptomyces* (van Dissel *et al.*, 2014). In the early stage of division, SsgA orchestrates division by facilitating the correct localization of SsgB. With the help of the transmembrane protein SepG, SsgB directly recruits FtsZ to the future septum sites and tethers the Z-ring to the inner membrane (Willemse *et al.*, 2011; Zhang *et al.*, 2016).

SsgB is the archetypical SALP, with a conserved function in the development of actinomycetes (Xu *et al.*, 2009). While the SsgB protein sequence varies strongly between less related Actinobacteria, the protein is extremely well conserved within a genus, with a maximum of one amino acid variation, a feature that has been applied for the phylogenetic analysis of closely related Actinobacteria (Girard *et al.*, 2013). In this work, we investigated the importance of individual residues in the localization of the septum during sporulation-specific division, by creating a library of SsgB mutants and studying their effect on cell division and morphogenesis. Single aa changes in SsgB had major effects on cell division, spore-wall synthesis, and DNA condensation and/or segregation. Remarkably, specific mutations led to the formation of additional septa with 10° to 90° rotation of the division plane. Such longitudinal fission has so far only been seen in the worm-associated bacteria *Candidatus Thiosymbion oneisti* and *Thiosymbion hypermnestrae*. In these two bacteria, cell growth and longitudinal division are polarized by their symbiotic nematode hosts (Pende *et al.*, 2018). X-ray crystallography revealed major structural differences between the SsgB from *S. coelicolor* and its distant orthologue from *T. fusca*. Our data support the predominant role of SsgB in the accurate positioning of the division site and the placement of the Z-ring.

## RESULTS

### Mutational analysis of SsgB

SsgB shows unusual conservation, with near complete conservation within a genus, and high divergence even between related actinobacterial genera. To investigate this further, we analyzed the effect of point mutations in SsgB on cell division and morphogenesis, using *S. coelicolor* as the model system. For this, we first created a random mutant library using error-prone PCR, similar to the approach used previously for the mutational analysis of SsgA (Traag *et al.*, 2007). Mutant genes, preceded by (and transcribed from) the original *ssgB* promoter region, were cloned into the low-copy number vector pHJL401 and introduced into the *ssgB* null mutant, followed by scrutiny of sporulation and cell division. To ascertain that the observed phenotypes were not due to differences in SsgB expression, the mutant was also complemented with a clone expressing wild-type SsgB, which gave a wild-type sporulation phenotype (see below). Additionally, Western analysis was performed using anti-SsgB antibodies. Samples were equalized for protein content and corrected based on the levels of elongation factor EF-Tu1 (Vijgenboom *et al.*, 1994). This revealed an average expression level of 77% +/- 10% of the wild-type level.

Spores of *S. coelicolor* are grey-pigmented due to the production of the WhiE spore pigment (Kelemen *et al.*, 1998), while colonies developing non-sporogenic aerial hyphae are white; intermediate phenotypes (reduced sporulation results in a light-grey pigmentation) also occur. This feature was utilized to subcategorize all transformants into three groups: white, light grey and grey. The mean grey level of growing patches was analyzed based on the scanner images. By this approach, the degree of sporulation could be readily monitored (Figure S1 and Table S1). 232 clones were isolated from the transformants and sequenced. Of these, 84 had no or silent mutations, 39 had multiple mutations and 65 had insertions or deletions. Of the 42 remaining clones, 35 unique single substitutions were identified and these were analyzed further. Out of 35 SsgB variants, six failed to sporulate and the others showed significant sporulation defects or reduced sporulation (Figure S2, Table S1).

To obtain more detailed insights into the morphological changes correlating to the substitution mutants, the transformants expressing SsgB variants were subjected to transmission electron microscopy (TEM) (Figure 1). Wild-type spores were homogeneous in size, with a thick electron-dense spore wall and condensed DNA in the centre of the spores. Conversely, spores from transformants expressing SsgB variants generally showed high variation in spore-wall thickness, spore size and shape, and frequently also aberrant DNA segregation and/or condensation (Figure 1). Much to our surprise, in some cases up to 90° rotation of the septal plane was seen, dividing the spores parallel to the growth direction of the hyphae. This suggests that mutation of single SsgB residues may affect the coordination of cell division in aerial hyphae of *Streptomyces*, as detailed below.

**Figure 1.**
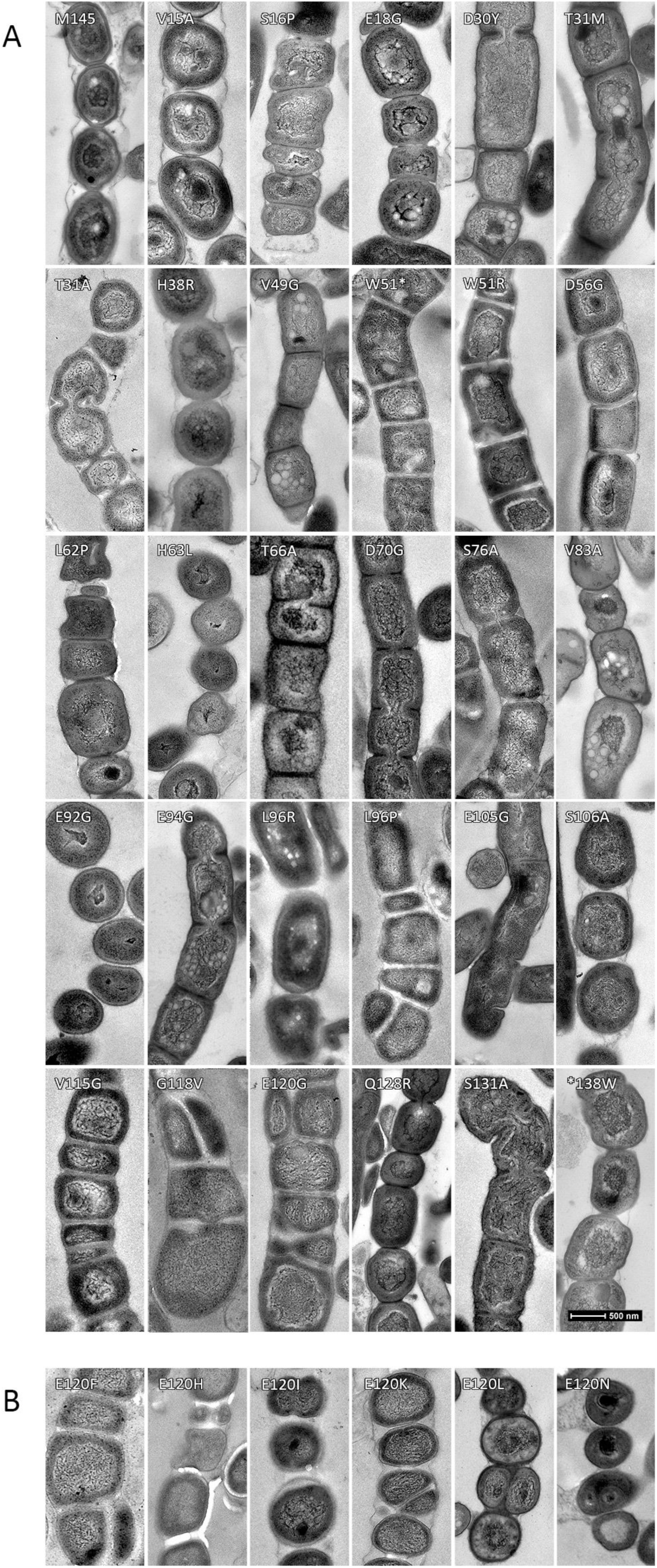
Transmission electron micrographs of sporogenic hyphae from the WT strain and from transformants expressing SsgB substitution mutants. (A) SsgB substitution mutants resulted in pleiotropic sporulation defects. T66A showed normal sporulation; D70G, S76A, E92G, S106A resulted in thinner cell walls; D56G, L88R, L96R and L96P affected DNA condensation and/or segregation; L96P, V115G, G118V, E120G gave rise to the formation of additional diagonal or longitudinal septa. V15A, S16P, E18G, D30Y, T31A, T31M, H38R, V49G, W51R, L62P, H63L, V83A, E94G, E105G, Q128R, S131A all showed highly variable spore sizes (see also Table S2). Bars: 500 nm for all TEM micrographs. (B) TEM images of spore chains from six additional transformants expressing substitution mutants of SsgB (E120F, H, I, K, L or N) that also gave rise to longitudinal division (for SsgB E120G see Fig. 1A).

### Rotation of the division plane due to single amino acid substitutions in SsgB

Based on the outcome of the random mutant library, 22 residues were selected for site-saturated mutagenesis, namely W51, L88, A95, L96, L97 and the C-terminal residues 115-131 that are centered around E120 that correlated to the surprising longitudinal division (Table S2). Each of these residues were changed into on average 14 different amino acid residues using DNA synthesis (Table S2). Mutants for the hydrophobic residues W51, L88, A95, L97, V115, P116 and P117, frequently had non-sporulating phenotypes (Table S2). Variable spore sizes were seen in most of the mutants, with some also showing irregular cell wall thickening (Figure 1). Importantly, thirteen mutants wherein E120 was replaced by either A, C, F, G, H, I, K, L, N, P, Q, S or T produced septa with significant rotation of the division plane - in addition to canonical septa. The angles of these ectopically positioned septa ranged from diagonal to longitudinal (i.e., 90° rotation, with septa parallel to the hyphal wall), of which 5.2 % were positioned diagonally (529 out of 10257), and 0.8 % longitudinally (86 of the 10257). See Figure 1B and Table S3. In addition to mutants expressing SsgB E120 variants, longitudinal division was also seen in mutants expressing SsgB variants V115G, G118V, L96I, L96P and L96S, whereby the latter three produced relatively few ectopic septa (Table S4). To the best of our knowledge, this is the first report of longitudinal cell division in any free-living bacterium. To ascertain that longitudinal division does not occur in the wild-type strain under the chosen conditions, over 1000 samples of the wild-type strain were checked by SEM and TEM, and not a single rotated septum was observed.

In order to see if the longitudinal septation also resulted in physical separation of the severed spores, we made impression prints of the strain expressing SsgB variant E120G, that had been grown for 7 days on SFM agar plates. These spores were then fixed with 1.5% glutaraldehyde in PBS, followed by dehydration using a graded series of acetone (70-100%). This experiment clearly demonstrated that the strain expressing SsgB E120G produces spores that are longitudinally sectioned in two and that this process is completed by spore fission (Figure 2A, panels e-f). The fixation procedure led to some drying artifacts in wild-type (Fig. 2A, panel a) and ssgB E120G (Fig. 2A, panels c-d) spores, but this effect was clearly different from the longitudinal division.

**Figure 2.**
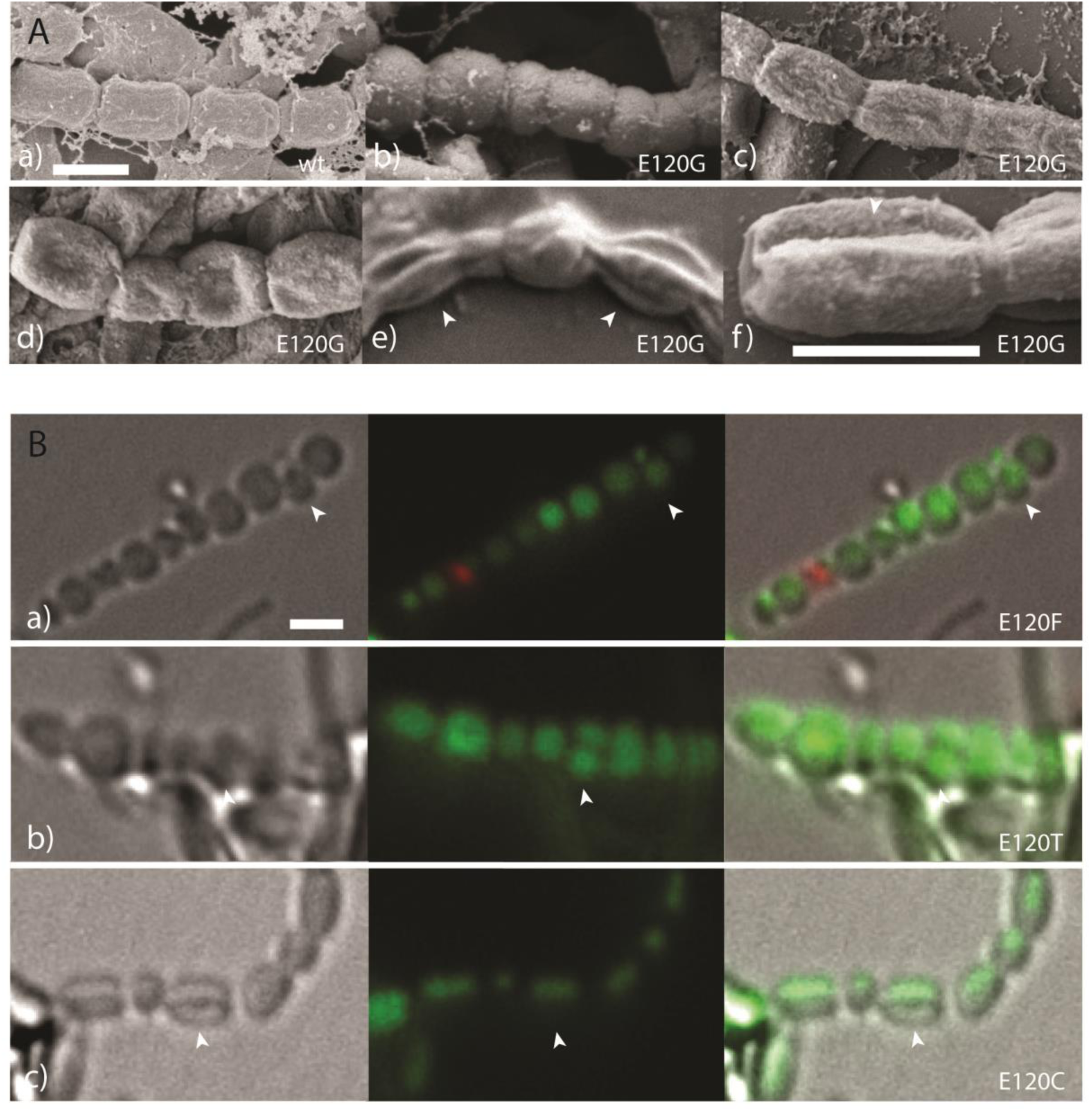
Impression prints of spores from cells expressing wild-type SsgB or SsgB E120 mutants. (A) SEM imaging. The fixation procedure led to occasional artefacts in the form of mild damage to spores (panels a, c) and sometimes to collapse of spores (panel d). However, the collapsed spores can clearly be discriminated from those that had been divided along the horizontal axis (panel e, f). (B) Longitudinal cell division (pointed by arrowhead) revealed by fluorescent microscope for different E120 mutants. *Left*, bright-field images; *middle*, Syto9/PI stained images; *right*, overlays of the two images. All the spores were obtained after 7 days of growth from wild-type cells or from transformants of its *ssgB* null mutant expressing SsgB E120 mutants. Bars, 1 μm.

Viability of the spores of mutants in which the residue E120 had been replaced by other amino acids were compared to those of wild-type spores. For this, impression prints were stained with Syto9 for viable spores and propidium iodide (PI) for dead spore, and imaged via fluorescence microscopy. While wild-type SsgB spores were nearly all viable, those obtained from E120 substitution mutants varied a lot, with 5-70% dead spores, depending on the mutant (Figure S3). Like in de SEM experiments, longitudinal septation could also be seen from the outside using light microscopy, indicative of unique cell fission parallel to the hyphal wall (Figure 2B, panels a-c).

### Localization and dynamics of SsgB variants

To confirm that longitudinal division in the aerial hyphae correlated to the localization of SsgB, chimeric SsgB-eGFP and SsgB-G118V-eGFP fusions were created as described (Willemse *et al.*, 2011). While wild-type SsgB (Figure S4A, panel a) showed the typical pattern of foci on either side of the hyphal wall, the G118V variants localized more centrally and also longitudinally (Figure S4A, panels b and d). This resulted in both canonical septal rings (perpendicular to the hyphal wall) and with a certain frequency also septa that were tilted by 90° (marked by arrowheads in Figure S4A, panel c).

Fluorescence intensities of the chimeric fusions were measured on the same width of the hyphae (Figure 3). The plotted graph of wild-type SsgB indicates its localization on either side of the hyphae wall. Whereas, Δ*ssgB*::*ssgB*(G118V) and Δ*ssgB*::*ssgB*(E120G) showed aberrant localization, which occasionally resulted in longitudinal septation, where SsgB localized in the middle of the hyphae, consistent with the observed longitudinal cell division. To gain insights into the dynamic association/dissociation of SsgB with the divisome, monomeric exchange was examined via Fluorescence Recovery After Photobleaching (FRAP). The recovery time after photobleaching was determined both on pre-septation foci as well as on septa. No difference in dynamics was seen between wild-type SsgB and its G118V variant. Both showed a recovery time of around 60 s (Figure S4B), which is similar to previously reported data (Willemse *et al.*, 2011).

**Figure 3.**
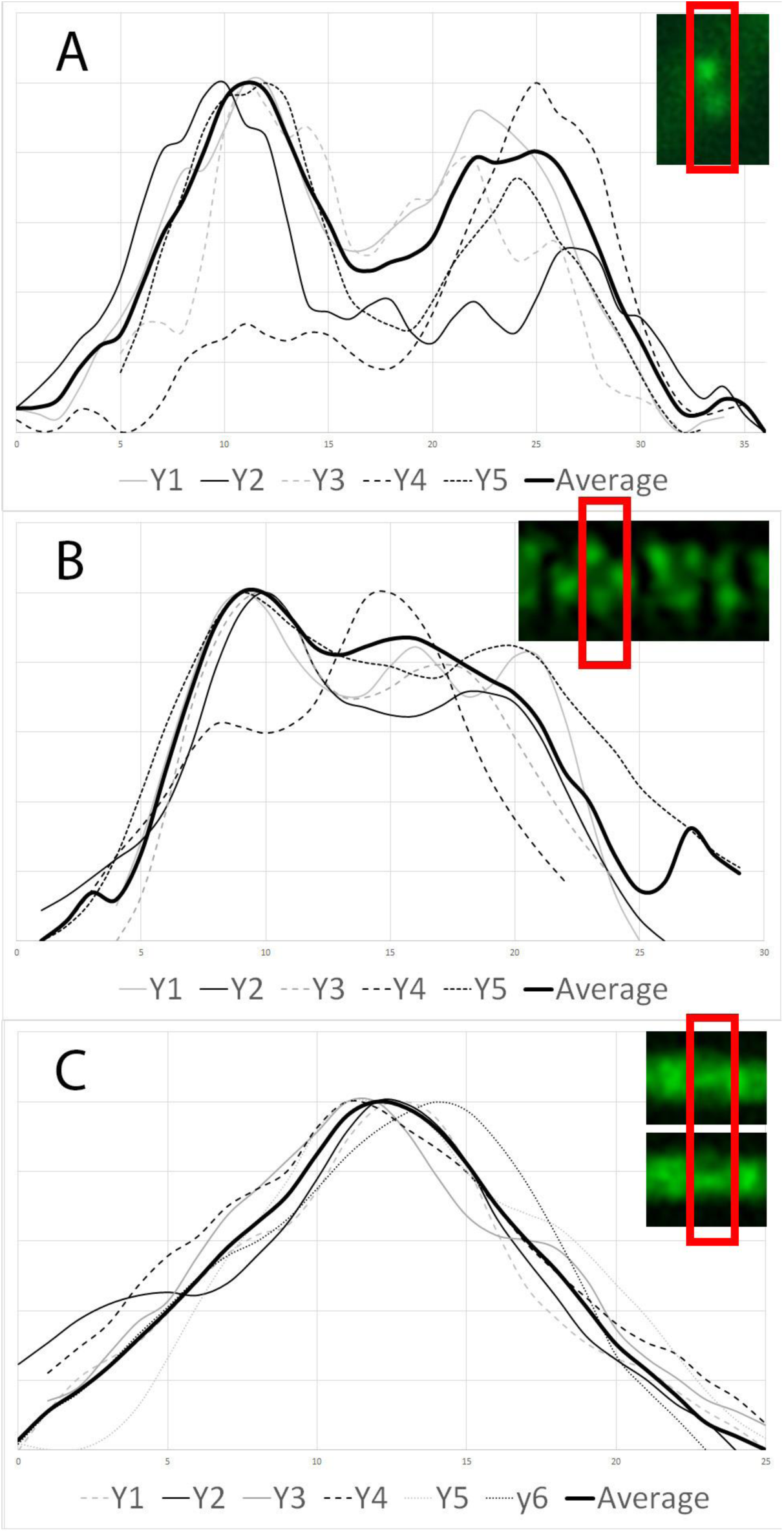
Intensity plots of SsgB foci on the septa. (A) The wild-type SsgB localization during early cell division shows two foci on either side of the hyphal wall; (B) Mutants expressing SsgB(G118V) or SsgB(E120G) showed aberrant localization, whereby SsgB was located all over the hyphal wall; (C) Occasional longitudinal septation was seen, whereby eGFP fusions of SsgB mutant proteins localized parallel to the hyphal wall and in the middle of the hyphae. The red box indicates 630 the width of the box that was used to produce the profiles.

DNA content of over 500 spores of wild-type strain and Δ*ssgB*::*ssgB*(E120G) was studied. While wild-type spores showed a normal DNA distribution, Δ*ssgB*::*ssgB*(E120G) showed major variation in DNA content (Figure S5A). As an illustration, one spore chain containing longitudinal divisions is shown with the respective DNA content in each spore, which revealed 0.4 to 3.0 chromosomes for each spore in Δ*ssgB*::*ssgB*(E120G) (Figure S5B). Conversely, 0.83 to 1.15 chromosomes were observed in the wild-type strains for each spore (Figure S5C).

### SsgB-E120 mutants assist in the polymerization of FtsZ filaments

To establish whether SsgB E120G and E120A had retained the ability to interact with FtsZ, wt SsgB, SsgB variants E120A and E120G, as well as a C-terminally truncated version of SsgB (SsgBΔC, spanning 1-114 aa) were expressed and purified and then tested using a pelleting assay. SsgB of *S. coelicolor* (*Sc*SsgB) was enriched in the pellet after incubation with FtsZ under polymerizing conditions (i.e. in the presence of GTP and Mg^2+^), while about 50% of FtsZ was recovered by centrifugation (Figure 4A). *Sc*SsgB variants E120A, E120G and SsgBΔC all pelleted in the presence of FtsZ when GTP and Mg^2+^ were added (Figure 4A), although less efficiently as compared to wt *Sc*SsgB. Neither FtsZ nor SsgB was recovered by centrifugation in the absence of GTP (Figure 4A), confirming that they did not form aggregates.

**Figure 4.**
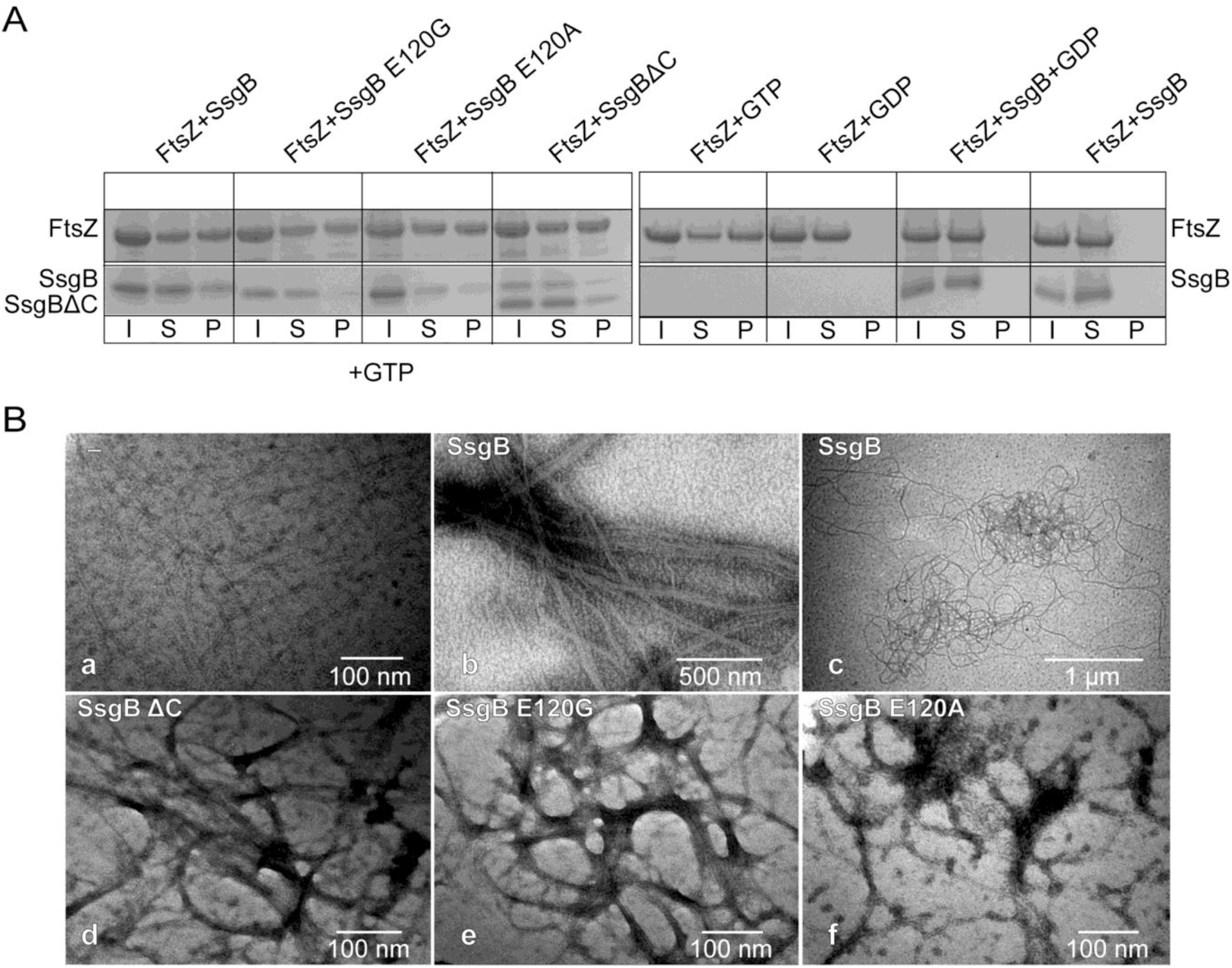
Co-localization and interactions between SsgB substitution mutants and FtsZ. (A) FtsZ sedimentation assay with different SsgB variants, in the presence of either GTP or GDP. Note that GTP is required for sedimentation. Initial samples (*I*) were used to assess the total protein content in each reaction. Soluble (*S*) and pelleted (*P*) fractions were separated by centrifugation at 45,000 rpm. (B) TEM images of FtsZ filament structures formed in the presence of GTP and different SsgB variants: (a) FtsZ alone; (b, c) FtsZ with wt SsgB; (d) FtsZ with SsgBΔC; (e) FtsZ with SsgB E120G; (f) FtsZ with SsgB E120A.

We then tested the polymerization of FtsZ by the same SsgB variants using negative staining. FtsZ alone formed short, straight and single-stranded filaments in the presence of GTP (Figure 4B, panel a). Addition of wt SsgB promoted the formation of bundled filaments (Figure 4B, panels b and c), similarly as seen for SsgB from *T. fusca* (Willemse *et al.*, 2011). Extended and bundled filaments were also observed in the presence of SsgB E120A, E120G and SsgBΔC, with E120A showing fewer and more ‘loose’ bundles (Figure 4B, panels d, e and f). Taken together, these data show that mutation of E120 or by deletion of the 23 C-terminal residues does not prevent the binding of *Sc*SsgB to FtsZ, whereby the mutant proteins still promote the formation of FtsZ filaments.

### Crystal structure of *S. coelicolor* SsgB

In order to gain more insights into the structure-function relationship for key SsgB residues, the structure of *Sc*SsgB was resolved via X-ray crystallography. For this, hexahedron crystals were obtained, and based on this, a homo-trimer was resolved at 2.1 Å (PDB ID Code 6SLC) with eight molecules per asymmetric unit (Table 1). The 13 aa residues at the C-terminus were highly mobile and could therefore not be modeled, due to lack of electron density. Each subunit was arranged as an α + β fold, with seven β-strands packed into a barrel structure, covered by three α-helices (Figure 5A), which strongly resemble those of *Tf*SsgB structure(Xu *et al.*, 2009) (Figure 5C). The root-mean-square deviation (r.m.s.d.) is 1.9 Å with 92% of all residues aligned in *Sc*SsgB and 87% in *Tf*SsgB (aa sequence identity between these two proteins is 46%). A superimposition of *Sc*SsgB and *Tf*SsgB subunits is shown in Figure 5E. *Sc*SsgB trimer adopts a “whirly” shape and is assembled through an antiparallel β-sheet interaction between β1 from one subunit and β4 from the neighbouring subunit (Figure 5B, Figure S6). In contrast, the *Tf*SsgB trimer is assembled mainly through α-helices (Figure 5D). The *Sc*SsgB trimer forms a 12-stranded beta-barrel with an inner diameter of about 20-25 Å (Figure 5B).

**Table 1.**
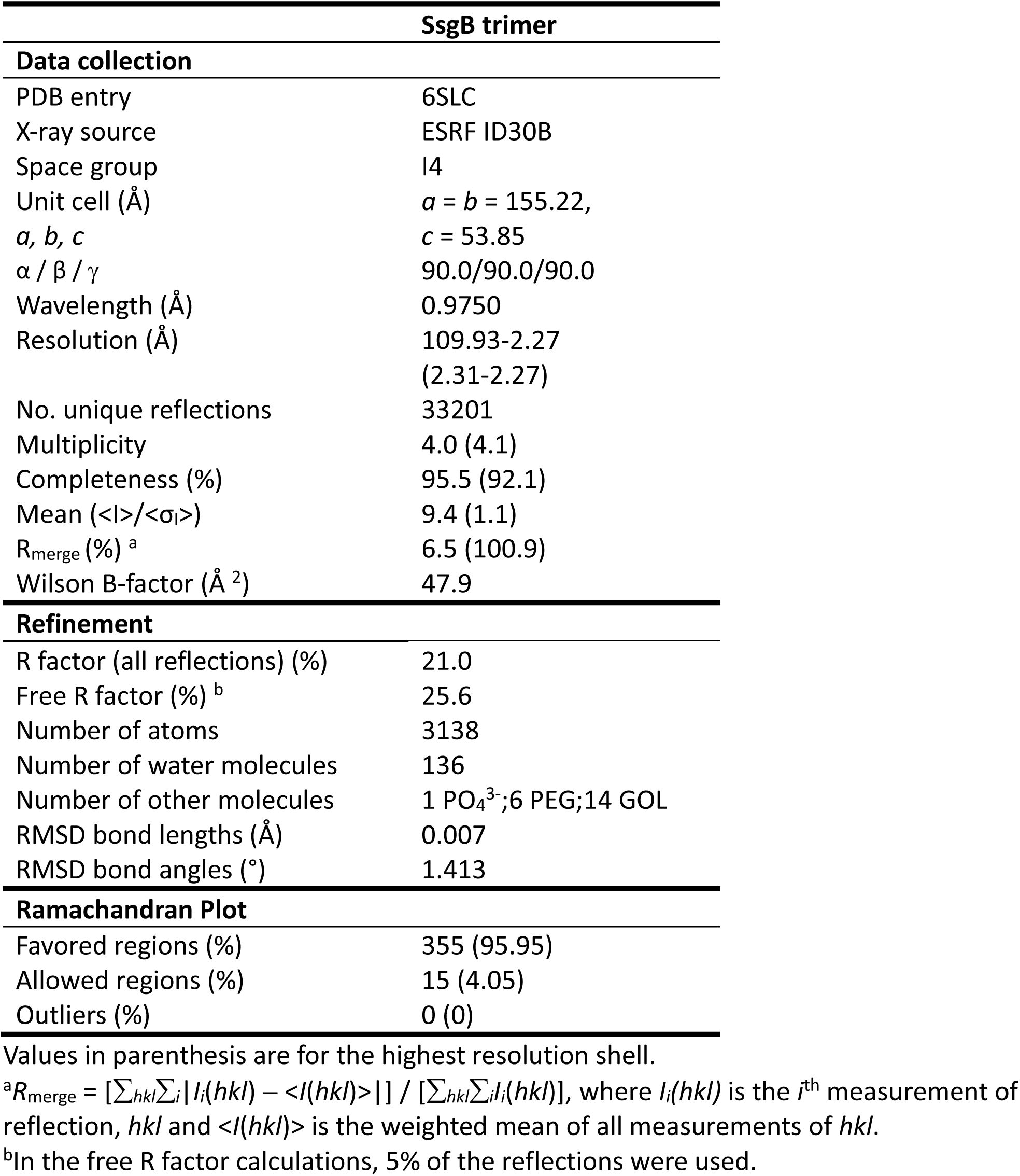
Data collection, model refinement, and final structure statistics.

**Figure 5.**
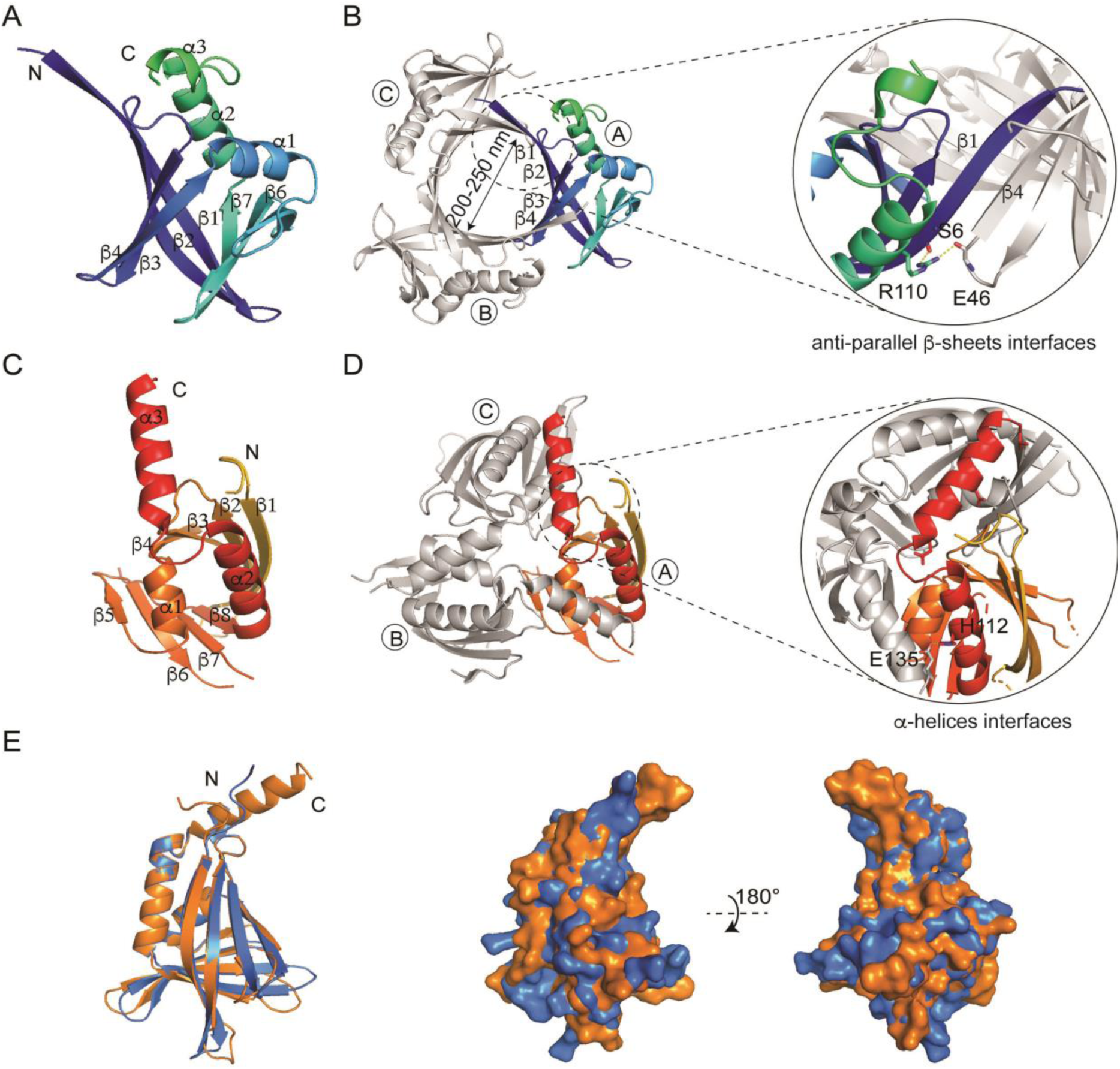
Crystal structure of the *Sc*SsgB trimer. (A) Ribbon diagrams showing the monomer structure of SsgB from *S. coelicolor*. (B) The overall structure of *Sc*SsgB reveals a trimer. Structure statistics are listed in Table 2. The interface between adjacent monomers is formed by two antiparallel β-sheets. (C) The monomer structure of SsgB from *T. fusca* (PDB code 3CM1). (D) The interface between adjacent monomers of *Tf*SsgB is formed by α-helices. (E) Overlap of *Sc*SsgB (blue) and *Tf*SsgB (orange) subunits. *Left*, side view of the electrostatic surface alignment of *Sc*SsgB and *Tf*SsgB structure. *Right*, the same electrostatic figure but rotated by 180°.

### Analysis of single mutations and mapping of key residues

Key residues that correlated to the occurrence of longitudinal cell division were mapped onto the *Sc*SsgB trimer structure (Figure 6A, Figure S7A). Mutations that correlate to residues that are evolutionary conserved in all SALPs are underlined (Figure 6A, Figure S7B). Residues V115, G118 and E120 cluster together, and are centered on the lid of the beta-barrel consisting of α1, α2-α3 and β1-β2 loop, which is close to the interface between α3 and the rest of the subunit (Figure 6B). Consistent with their strategic localization in the α2-α3 loop, amino acid substitutions V115G, G118V or E120G resulted in the formation of aberrant tilted septa in addition to the canonical septa perpendicular to the hyphal wall, with some septa showing full 90° rotation of the division plane. Residue E120 plays a key role in maintaining the proper angle between α3 and the rest of the protein. Three hydrogen bonds are formed between the E120 side chain and the main chains of E120, T119 and G118 in the α2-α3 loop region. Besides, E120 and V115 provide two additional salt bridges to R55. Interestingly, a π-π interaction and a hydrogen bond were observed for E120-Y35 and H121-Y35. All these interactions stabilize the angle of α3 (Figure 6C), supporting the importance of the proper angle between α3 and the rest of the protein to the function of *Sc*SsgB.

**Figure 6.**
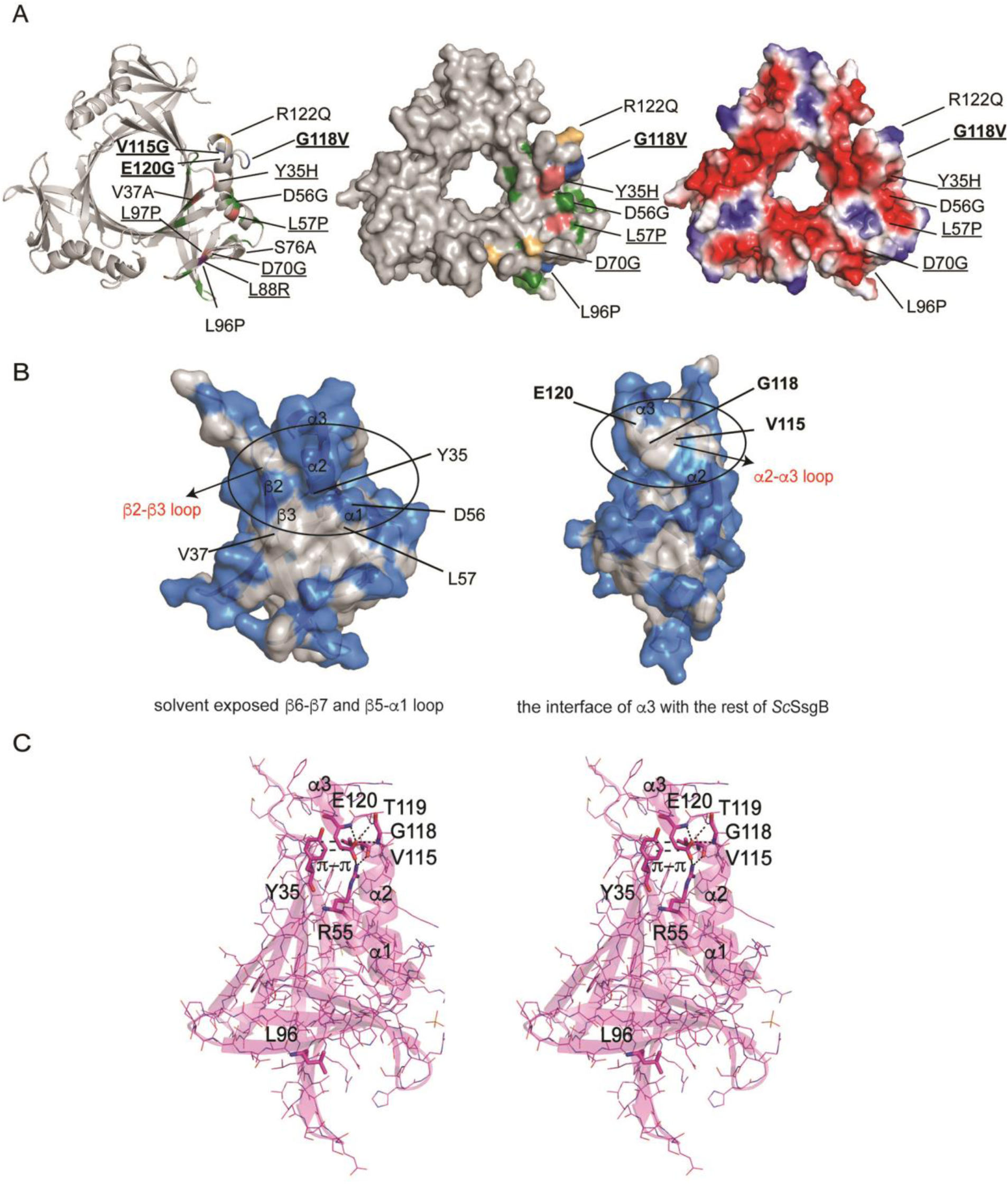
Key mutations and their interactions in SsgB trimer structure. (A) *Left*, mutant residues L96, <u>G118</u>, <u>E120</u> (marine) which showed tilted division are highlighted; Mutation of residues <u>Y35</u>, V37, <u>L57</u>, <u>L97</u> (deep salmon) resulted in a sporulation block; mutation of residues <u>D70</u>, S76, R122 (light orange) led to thinner cell walls; variants D56, L96 and <u>L88</u> (violet purple) had damaged DNA. *Middle, Right*, top view of all the functional mutants mapped on the surface structures. Conserved residues are underlined. (B) Key mutations (V115, G118 and E120) are clustered on the lid of the β-barrel, consisting of α1, α2-α3 and β1-β2 loops. (C) Stereo view of E120 in the monomer structure and its interactions with the surrounding residues.

### Molecular dynamic simulation of SsgB E120G

Our work demonstrated that residue E120 plays a key role in maintaining the proper angle of α3 relative to the rest of the protein. Mutation of this residue would disrupt the interaction, and changing the angle of α3 may drive rotation of the septum plane, and thus explain the observed longitudinal cell division. The heading and the high B-factor of the existing residues in the α3-helix (Figure S8) suggests that it can extend flexibly to the center of the trimer and serve as a lid for the mentioned beta-barrel at the center of the structure. Despite many attempts under different crystallization conditions, we failed to obtain crystals for SsgB E120G or SsgB E120A. In silico molecular dynamics revealed that while the α3 helix of the wild-type SsgB protein is not affected significantly by the simulations, with a 2.7 and 3.0 Å of distance respectively between E120 and R55 (Figure 7A), the α3 helix of SsgB E120G flips some 90° outwards of the tight trimer, with the distance increasing to 10.6 and 10.9 Å, respectively (Figure 7B). This provides supportive evidence that the angle of the α3 helix may indeed play a key role in determining the orientation of the septum plane.

**Figure 7.**
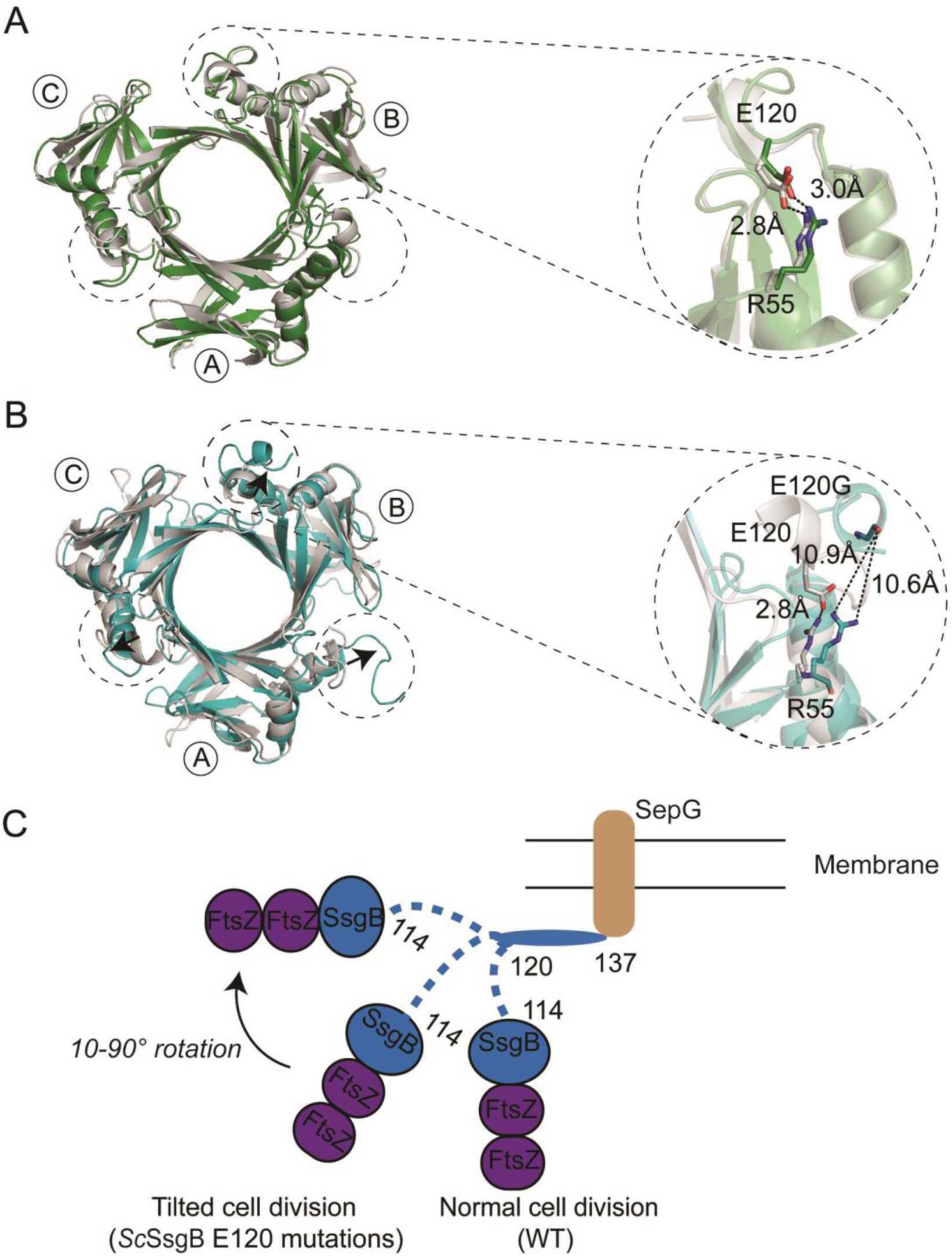
Molecular simulation of SsgB wt and E120G mutant. (A) MD result of SsgB wt structure. The α3 helix stays at the same orientation and the distance between R55 and E120 keeps at 2.8 Å and 3.0 Å, before (green) and after MD (grey). (B) The arrows indicate the changed angle of SsgB E120G mutant (cyan) compared to the wild type SsgB (grey) after MD. (C) Model for how E120G mutant enhances longitudinal cell division.

### Oligomerization Studies of SsgB

Crystallographic data obtained for SsgB from *T. fusca* (Xu *et al.*, 2009) and for *S. coelicolor* (this work) suggested that SsgB forms trimers. To ascertain this, size-exclusion chromatography (SEC) of wt *Sc*SsgB and *Sc*SsgBΔC was conducted. SEC results indicated that freshly purified wt *Sc*SsgB and *Sc*SsgBΔC mainly existed as a monomer in solution (Figure S9A), while the SEC experiment of the same batch of protein showed the existence of a trimer after a short-time storage in −80°C, revealing the conversion of monomers to trimers over time (Figure S9B).

## DISCUSSION

SALPs play a central role in controlling the steps of sporulation-specific cell division in *Streptomyces*. Inspired by the extremely high conservation of the SsgB protein in *Streptomyces* species, with natural variants only found in aa 128 (Q, R or T), we studied the effect of point mutations in the protein on cell division and morphogenesis of the model strain *S. coelicolor*. As expected, many mutants showed morphological defects relating to cell division and sporulation, including varying spore sizes, aberrant DNA segregation and condensation, and cell wall thickening. Surprisingly, SsgB substitutions L96P, V115G, G118V and various changes in E120 caused the formation of additional septa that section the spores diagonally or longitudinally, perpendicular to the canonical septa. This is the first example of longitudinal division in a free-living bacterium. Diagonal and longitudinal Z-rings and septa always coincided with canonically oriented Z-rings/septa. Furthermore, the longitudinal Z-rings connect two canonical Z-rings. And finally, the longitudinal septa could be formed with different spacing relative to the hyphal wall, allowing asymmetric cleavage of spores. This strongly suggests that during normal Z-ring formation, a second Z-ring is formed under different angles ranging from 45-90 degrees. Eventually, the longitudinal septation also resulted in physical separation of spores along the horizontal axis, as seen by SEM and light microscopy. This shows that these ectopic cell division events were completed via cytokinesis. We previously showed that enhanced expression of SsgA, a cell division activator that assists in the localization of SsgB, results in enhanced cell division and even the formation of ectopic spores in vegetative hyphae (van Wezel *et al.*, 2000). This underlines that SsgA and SsgB play a pivotal role in determining where septa are positioned in the hyphae of streptomycetes.

Mutational and structural analysis, fluorescence imaging and 271 molecular dynamic simulation of *Sc*SsgB provided more insights into the structural basis for the observed longitudinal cell division. The hydrogen bonds (E120-E120, E120-T119 and E120-G118, H121-Y35), salt bridges (E120-R55 and V115-R55) and π-π interaction (E120-Y35) stabilize and maintain the proper angle of the α3 helix. Substitutions in E120 disrupts this critical interaction, and most likely results in major changes in the orientation of α3, by up to 90 degrees. Mutations of the interacting partners all showed functional defects, as seen from the blocked cell division (Y35H), septum rotation (V115G, G118A), DNA segregation (V115G, G118A) and heterogeneity in spore sizes (V115G, G118A, H121I). Moreover, some longitudinal septa were seen in L96P mutants, which again can be explained by changes in the orientation of helix α3, as the mutation will disrupt the interaction between the neighbouring α2 helix and β7-strand. We therefore propose that rotation of helix α3 is the driving factor for rotation of the Z-ring for up to 90 degrees along the long hyphal axis, corresponding to the rotation of the septal plane seen in various mutants (Figure 7C). We propose the following order of events leading to longitudinal cell division: (1) SsgB localizes to the septum sites and recruits FtsZ, thereby assists in its polymerization and Z-ring formation; (2) in strains expressing specific SsgB variants (particularly in G118, E120 and V96), an additional second Z-ring is formed perpendicular to the canonical Z-rings, re-orienting the divisome to the central septum plane, parallel to the long axis of the hyphae. This eventually results in diagonally or horizontally severed spores. SsgB G118V, E120G and E120A had retained the ability to promote the assembly of FtsZ filaments in vitro. FRAP further confirmed the *in vivo* studies by showing that the canonically localized foci and the centrally localized foci produced in SsgB substitution mutants have the same dynamics, and that these are similar to those of wild-type SsgB.

So far, longitudinal fission had only been reported in studies by Bulgharesi and colleagues on the nematode-associated gammaproteobacteria *Candidatus Thiosymbion oneisti* and *Thiosymbion hypermnestrae* (Pende *et al.*, 2018). Longitudinal cell division in these bacteria is host-polarized by their nematode symbionts. The symbionts grow along the long axis and with increased cell width. The machineries for growth and division are not reoriented; instead, they mesh 295 to a point where they appear as “squeezed” *E. coli* cells (Pende *et al.*, 2018). Our work shows that also in free-living bacteria, and specifically in *Streptomyces*, cell division along the longitudinal axis is possible, caused by single aa substitutions in the FtsZ-recruiting SsgB. It will be interesting to see if longitudinal division can also be achieved in other bacteria, for example in bacteria where cell division is also positively controlled, such as in *Myxococcus xanthus*. In *Myxococcus*, the ParA-like ATPase PomZ recruits FtsZ to midcell during vegetative growth (Treuner-Lange *et al.*, 2013). While this system is different from that controlled by SsgB, it will be worth investigating whether amino acid substitutions in PomZ may achieve similar changes in Z-ring positioning.

In conclusion, our work shows that specific residues in SsgB, and especially residues G118 and E120, play a key role in stabilizing the SsgB structure. Mutations in these residues result in major changes in the control of Z-ring synthesis, resulting in additional septa that are formed diagonally or perpendicular to the canonical septa, thereby severing spores in two halves. This underlines the crucial role of SsgB in cell division control in streptomycetes.

## MATERIALS AND METHODS

### Strains and culturing conditions

All strains described in the paper are listed in Table S6. *S. coelicolor* M145 was obtained from the John Innes centre strain collection. Its *ssgB* null mutant was published previously (Keijser *et al.*, 2003). Transformants harbouring SsgB-expression vectors based on the low-copy number shuttle vector pHJL401 (Larson and Hershberger, 1986) were grown on SFM agar plates containing 50 μg/ml apramycin (for the *ssgB* deletion) and 25 μg/ml thiostrepton (to maintain the plasmid) at 30°C. For growth in liquid medium the recombinants were grown in a 1:1 mix of TSBS and YEME at 300C. *E. coli* JM109 was used for amplification of plasmids and *E. coli* Rosetta™ 2 (DE3) pLysS for overexpression and isolation of the His_6_- tagged proteins.

### Random mutagenic PCR

The SsgB promotor region and structural gene were amplified separately from the *S. coelicolor* chromosome using oligonucleotide pairs pSsgB_fw + pSsgB_rv and SsgB_fw + SsgB_rv, respectively (Table S7). An NdeI restriction site was introduced overlapping the translational start codon to enable ligation of these fragments after mutagenic PCR. The *ssgB* gene was cloned into a variant of pUC19 wherein the unique NdeI site had been removed, thereby creating pJPM1. Construct pJPM2 was based on the low-copy number shuttle vector pHJL401 in which the original NdeI site had been removed, and contained the PCR-amplified *ssgB* promotor fragment (cloned as EcoRI-HindIII fragment). Mutations in *S. coelicolor ssgB* were introduced by random mutagenic PCR using pJPM1 as template, as described (Traag *et al.*, 2007). The mixture of mutagenized *ssgB* genes produced by error-prone PCR was then ligated as NdeI-HindIII fragments behind the natural *ssgB* promoter in pJPM2. The DNA was subsequently transformed into protoplasts of the *S. coelicolor ssgB* null mutant, thereby generating a collection of *Streptomyces* colonies each expressing a variant of SsgB from the natural *ssgB* promoter. Mutations were verified by DNA sequencing.

### Scanner based imaging

Plates were incubated at 30°C on a flat-bed CCD scanner and imaged every 30 min for 7 days. Automated scanning was performed using Quickscan (www.burrotech.com) activated by the windows task scheduler. Hereafter the image stack was analyzed for gray values using imageJ/FIJI. This was achieved by drawing an equal sized circle in the middle of the grown colonies and measuring the grey level intensity of the stack via the Measure stack plugin of ImageJ.

### Automated spore measurements

For each image in the folder the scale is set to correspond to the microscopes settings, hereafter the image size is increased to optimize for averaging of pixels values at later stages of the macro. After increasing the image size by 5% in 20 consecutive operations the edge detection filter of imageJ is applied. Everything above the default threshold is defined as a proper edge. The holes are filled and the spores and hyphae that are detected in this manner are defined as “in focus”. The original file is thresholded with default settings and combined with an AND operation with the sharp spores. Particles that fall within the range of spores are analyzed, with minimum size of 0.65 μm^2^, and a roundness value between 0.75 and 1.

### Microscopy

#### Live/dead staining and DNA quantification

For live/dead staining *Streptomyces* strains were grown on SFM agar plates, and after 7 days cover slips were pressed onto the colony and mounted in PBS containing 10 µM syto 9 and 10 μg·ml^-1^ Propidium iodide t. Fluorescence and light microscopy were performed as described previously (Willemse and van Wezel, 2009).

For DNA quantification, *Streptomyces* colonies were grown against coverslips (at 45° angle) on SFM agar, and after 7 days taken out of the agar samples were fixed with 2% paraformaldehyde for 5 minutes and washed with 70% ethanol. Subsequently spores were stained with 1 µM Syto green (ThermoFischer) and imaged with a Zeiss Axioplan 2, with 470/40 excitation and 525/50 emission. For localization studies, cover slips were immediately imaged with either the same microscope was used for imaging the SsgB-localization. For DNA quantification the total intensity of each separate spore in a spore chain was measured, to circumvent staining variation the median value of each spore chain was set to 1 to normalize the data. To have enough data to normalize spore chains that were measured consisted of a minimum of 10 spores. All images were background-corrected, setting the signal outside the hyphae to 0 to obtain a sufficiently dark background. Images were processed using Adobe Photoshop CS4 and FIJI.

#### Fluorescence recovery after photobleaching (FRAP)

FRAP was performed with a Zeiss Imager LSM 510, using 488 nm excitation and 505-550 nm detection as described (Willemse *et al.*, 2011).

#### Electron Microscopy

Cryo-scanning electron microscopy (cryo-SEM) was performed as described (Piette *et al.*, 2005). For SEM imaging of individual spores, impression prints of 7-day old confluent plates were obtained and fixed with 1.5% glutaraldehyde in PBS. After 15 min fixation, samples were dehydrated using a graded series of acetone (70%, 80%, 90%, 96%, 100%, 15 min each) and subsequently critical point dried. Before examination 10 nm Platinum/Palladium was sputter coated on the sample to prevent charging during imaging. All images were obtained with a Jeol 7600 at 5kV and a working distance of 8 mm.

For transmission electron microscopy (TEM), small cubes of colonies were fixed with 1.5% glutaraldehyde in PBS for 1 hour, postfixed with osmium tetroxide (1%) for 1 hour, and dehydrated with a graded ethanol series (70%, 80%, 90%, 100% 15 minutes each). Ultrathin sections of 70 nm were cut and examined using a Jeol 1010. Purified FtsZ (15 μM) was combined with an equimolar amount of wt SsgB, or one of its mutants (SsgB E120G, SsgB E120A and SsgBΔC) in a reaction buffer of 20 mM HEPES, pH 7.5, 150 mM KCl, 2.5 mM MgCl_2_. Polymerization was initiated by the addition of 2 mM GTP to the assembly reaction. 50 μl reaction mixture was incubated for 2 minutes at 37°C. 15 μl aliquot was placed on a carbon coated copper grid and negatively stained with 2% uranyl acetate for 10 minutes, then washed and dried. Images were collected using transmission electron microscope (Jeol 1010) operated at 70 kV with 670 pA·cm^-2^ density and recorded on a camera.

### Co-pelleting assay

For pelleting experiments, purified FtsZ, with or without equimolar ratios of wt SsgB, or its mutants (SsgB E120G, SsgB E120A and SsgBΔC) were mixed at a final concentration of 15 μM. The samples were pre-spun at 45 k r.p.m in a Beckman 40.1 rotor for 30 minutes. Then supernatants were transferred to new tubes and 2 mM MgCl_2_ and GTP were added. After 10 min of incubation at 30°C, the tubes were centrifuged at 45 k r.p.m at 20°C for 30 minutes. the supernatants were removed for analysis and pellets were washed with the same buffer aside from the proteins. Pellets were dissolved in SDS gel loading buffer. All the samples were analyzed on a 4%-16% Pre-cast SDS-PAGE gel (BIO-RAD).

### Protein expression and purification

For expression of His_6_-tagged fusions of SsgB and FtsZ in *E. coli*, the entire coding region of *ssgB* or its mutants were cloned as NdeI-HindIII fragments into pET28a (Novagen). Sequences of *S. coelicolor ftsZ* were codon optimized prior to cloning into pET28a. SsgB was overexpressed in *E. coli* Rosetta™ 2 (DE3) pLysS strain as a 157 aa-long fusion protein containing the 137 aa native polypeptide and an N-terminal tag, GSSHHHHHHSSG. SsgB point mutants E120G and E120A, or its deletion mutant (ΔSsgB, 1-114 aa; lacking the C-terminal 11 aa), were expressed in the same way. FtsZ was produced as C-terminal His-tag fusion. His_6_-tagged FtsZ proteins were purified using routine methods as described (Mahr *et al.*, 2000).

### Oligomerization studies of SsgB

The polymerization state of SsgB was analyzed by size-exclusion chromatography on an ÄKTA-pure system using column Superdex™ 75 Increase 10/300 GL and Superdex™ 75 Increase 5/150 GL at room temperature, respectively. The elution solvent was 20 mM HEPES, 150 mM KCl, 2.5 mM MgCl_2_, 5% Glycerol, 1 mM DTT, at pH 7.5, and the flow rate was 0.1 ml·min^-1^. Absorbance at 280 nm was monitored. The γ-globulin, conalbumin, ovalbumin, myoglobin, and vitamin B12 were used as molecular weight standards.

### Crystallization and structure determination

SsgB trimer crystals were grown at 18°C by using the sitting-drop vapor-diffusion method. In a typical experiment, 1 μl of the protein stock (6.2 mg·ml^-1^ protein) was mixed with 1 μl of a reservoir buffer consisting of 0.2 M Sodium chloride, 0.1 M Sodium/potassium phosphate (pH 6.2), 50% PEG200. Crystals were moved to the same condition supplemented with 20% glycerol and flash-frozen in liquid nitrogen.

Data were collected at beam line ID30B (McCarthy *et al.*, 2018) at ESRF (Grenoble, France, 2017). Images were collected with a 0.15° oscillation angle and an exposure time of 0.037 s per frame at 100 K. Crystals forms diffracted to 2.1 Å for SsgB trimer. The data was processed with XDS (Kabsch, 2010) and scaled using AIMLESS (Evans and Murshudov, 2013) from CCP4 package(Bailey, 1994). Phases of SsgB trimer were calculated by molecular replacement with SsgB^*Tfus*^ PDB entry 1C3M(Xu *et al.*, 2009) as a model using MOLREP (Vagin and Teplyakov, 1997). The structures were finalized by manual building in COOT (Emsley *et al.*, 2010) and refined with REFMAC (Murshudov *et al.*, 1997). Residues were in the most favored regions of the Ramachandran plot (Ramachandran *et al.*, 1963) as determined by PROCHECK (Laskowski *et al.*, 1993). Crystallographic data are summarized in Table 2. SsgB trimer structure was deposited in the PDB with code 6SLD, respectively.

### Molecular simulation

All atom molecular dynamics (AAMD) simulation of SsgB wt and SsgB E120G were prepared and run using Gromacs 2016 package (Hess *et al.*, 2008). The simulated protein, wide type or mutant type, was centered in a cubic box surrounded by water molecules and counter ions Na^+^ were added to keep the total charge of the simulation box to be zero. The bonded and non-bonded parameters were obtained from AMBER99SB force filed (Hornak *et al.*, 2006). The SPCE (Berendsen *et al.*, 1987) water model was used. The Particle-mesh Ewald method was used to treat the long-range electrostatic interactions (Darden *et al.*, 1993). A cutoff of 12 Å was used for non-bonded interactions. Temperature was maintained at 300 K using v-rescale thermostat (Bussi *et al.*, 2009); Pressure was maintained at 1 bar Parrinello–Rahman barostat (Parrinello and Rahman, 1981; Rahman and Stilling.Fh, 1971). The simulation box was cubic of length around 9.2 nm, with periodic boundary conditions applied to all dimensions. All simulated systems were stabilized though energy minimization, short NVT (10 ps) and short NPT (5 ns) MD simulations to relax to favorable conformations. NPT simulations at 300 K and 1.0 bar were performed upon the stabilized structures and 50 ns trajectories were collected for our analysis.

## Supporting information

All Supplemental Tables and Figures

## ACKNOWLEDGEMENTS

We gratefully thank to the European Synchrotron Radiation Facility, Grenoble, France. This work was supported by a grant from the Chinese Scholarship Council (CSC) to X.X.

## CONFLICT OF INTERESTS STATEMENT

The authors declare no conflict of interests.

