## Supplementary material for "Longitudinal cell division is associated with single mutations in the FtsZ-recruiting SsgB in *Streptomyces*": All Supplemental Tables and Figures

belonging to the manuscript

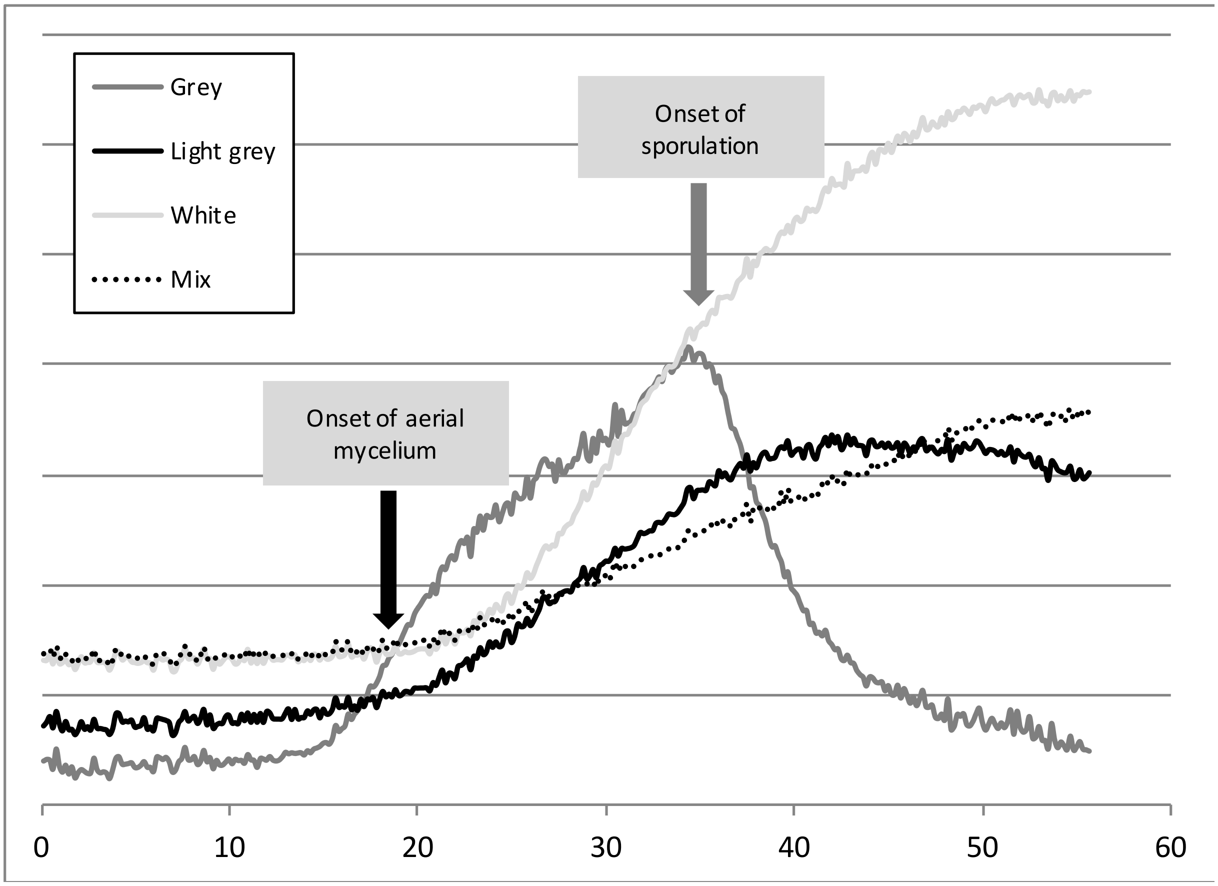


**Figure S1. Scanner-based analysis of pigmentation of *S. coelicolor* M145 expressing SsgB substitution mutants.** The degree of sporulation is detected by measuring the grey values of the strains over-time. The horizontal axis shows the time of inoculation in hours. Imaging was commenced 24 h post-inoculation.

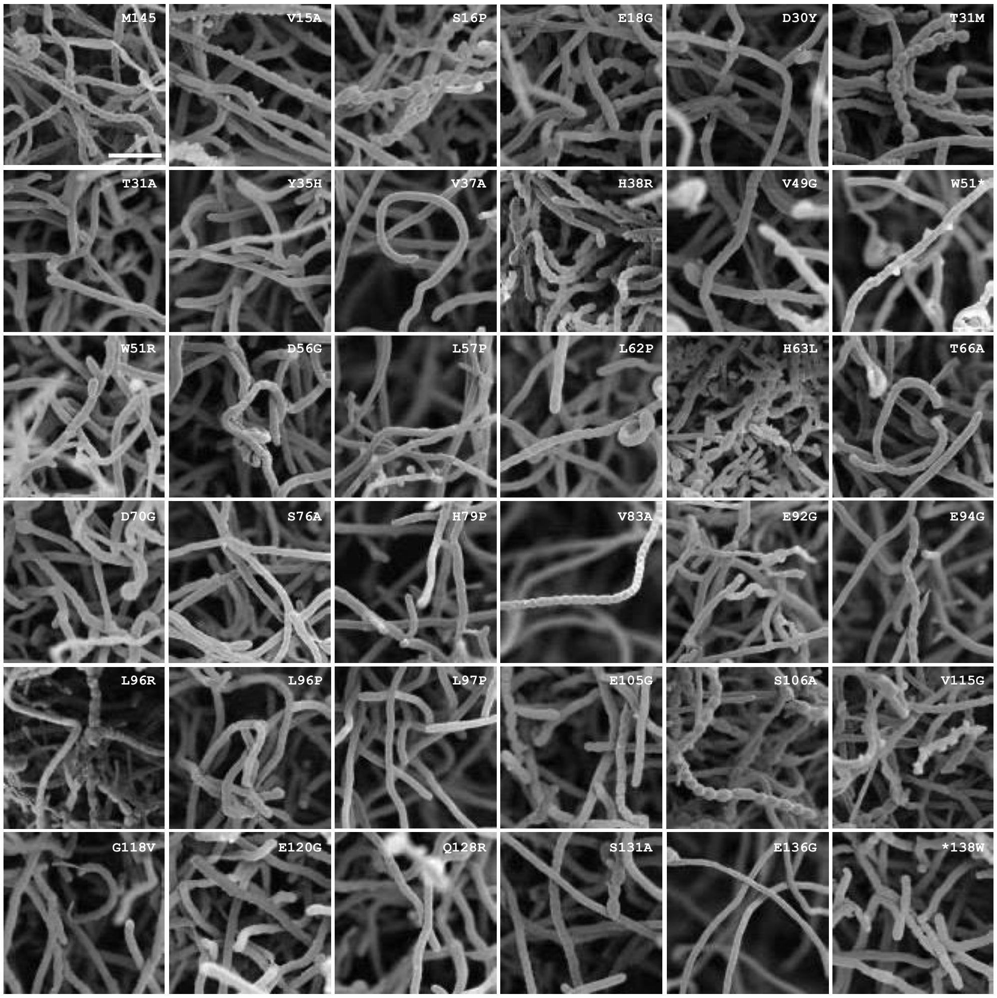


**Figure S2. Scanning electron micrographs of *S. coelicolor* M145 expressing different SsgB substitution mutants.** While most substitutions resulted in aberrant sporulation with defects in spore morphology, mutants Y35H, V37A, L57P, H79P, L97P, and E136G failed to sporulate (see also Table 1).

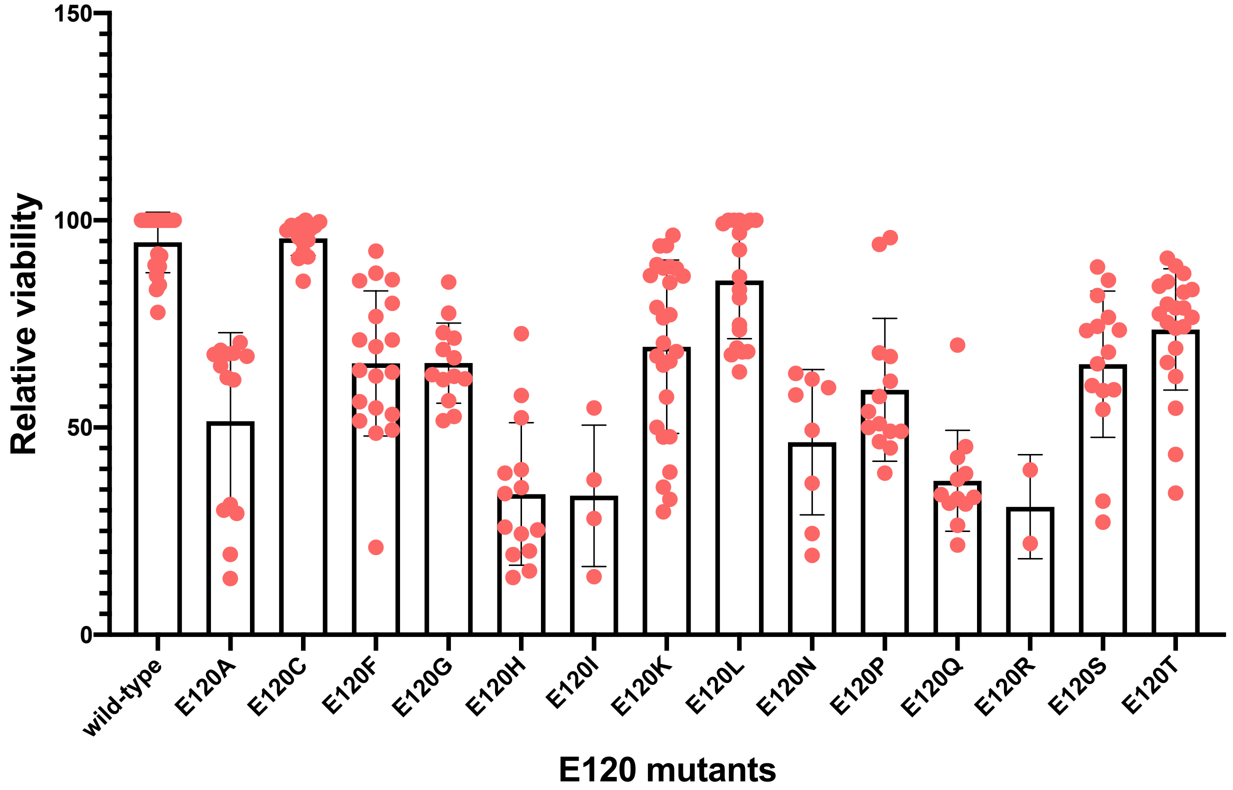


**Figure S3**. **Viability studies of spores from transformants of *S. coelicolor* M145 expressing various E120 mutants by Syto9 and propidium iodide staining.** Fluorescence micrographs were obtained for each individual E120 mutant and live/dead ratios calculated. Wild-type cells showed 95 - 100% viability, while viability of E120 mutant spores varied from 30 to 95%.

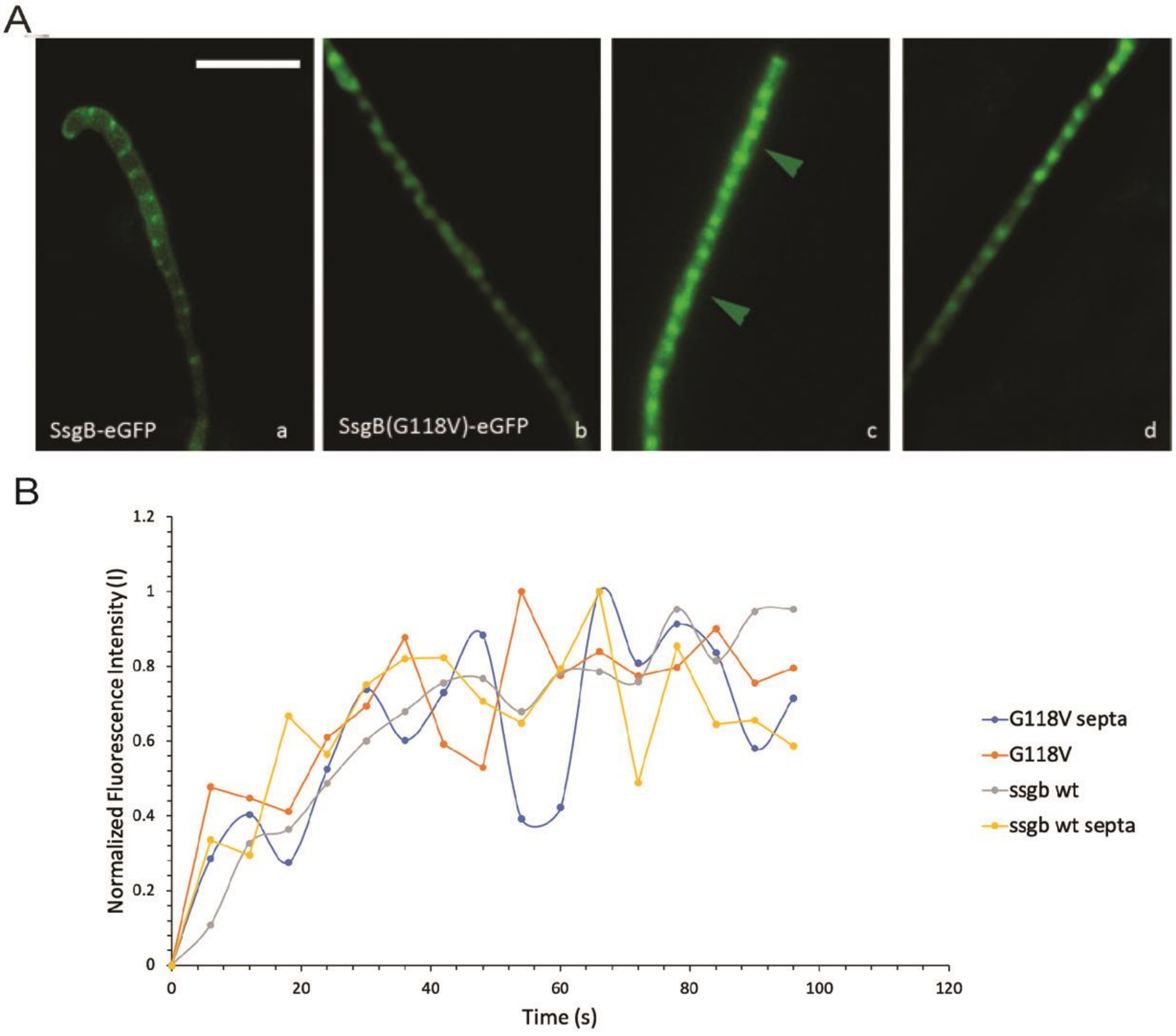


**Figure S4. Localization of SsgB *in vivo*.** (A) Subcellular localization of SsgB-eGFP fusion proteins. Wild-type SsgB-eGFP (a) had the typical pattern of foci on either side of the hyphal wall. The SsgB(G118V)-eGFP (b-d) protein regularly showed aberrant localization, including longitudinal cell division (arrowhead in c). Bar, 5 µm. (B) The association/dissociation time of wild type SsgB and its variant G118V showed by Fluorescence Recovery After Photobleaching (FRAP).

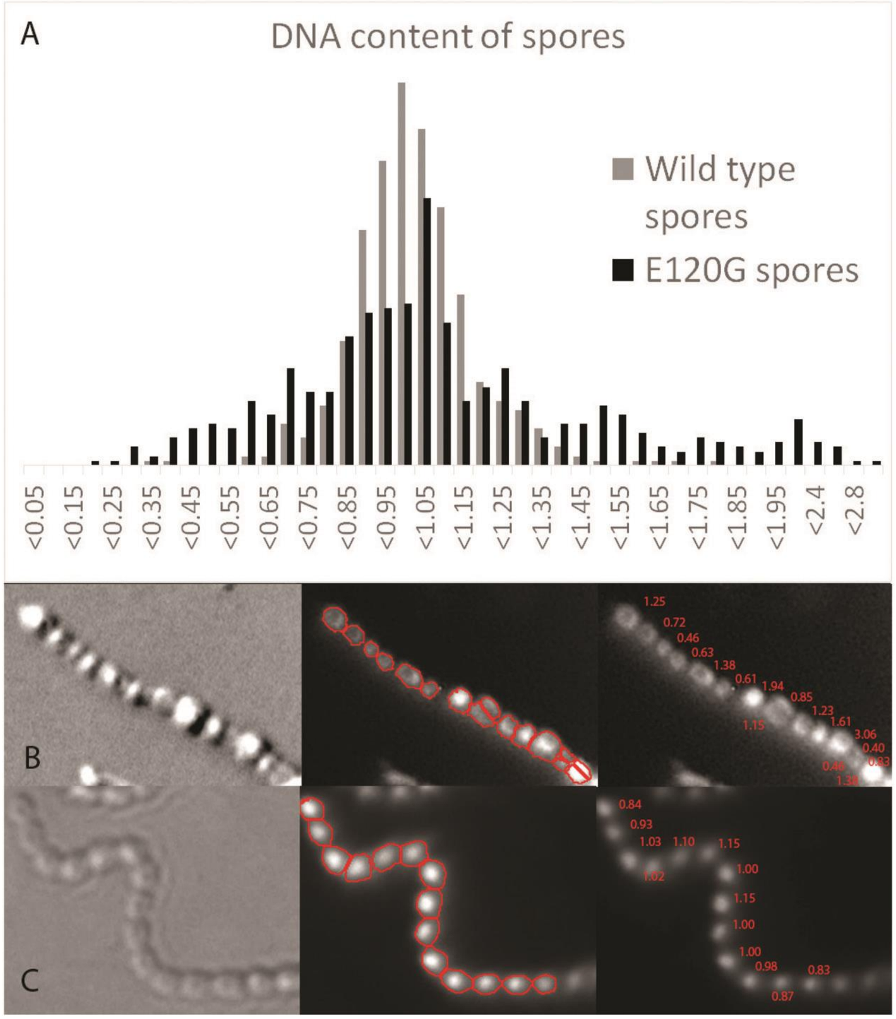


**Figure S5. DNA content analysis of spores of *S. coelicolor* M145 and its derivative expressing SsgBE120G.** (A) A normal distribution was observed for wild-type strain, whereas the strain expressing SsgB(E120G) had much more variation in DNA content. The median DNA content of the spores was set to 1 to internally normalize the data. (B) A representative spore chain containing longitudinal divisions is shown with the respective DNA content in each spore (between 0.4 and 3.0 chromosomes per spore). (C) In comparison, wt spores showed relatively little variation (between 0.83 and 1.15 in the spore chain).

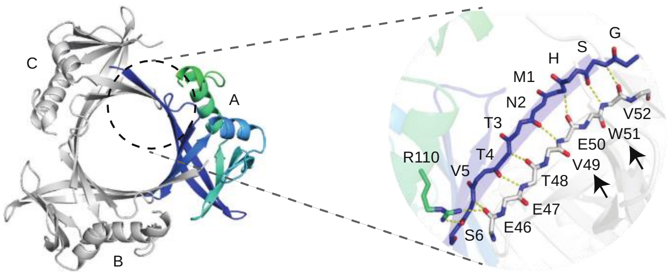


**Figure S6. Interfaces of the ScSsgB trimer structure.** Arrows indicate the key residues W51 and V49.

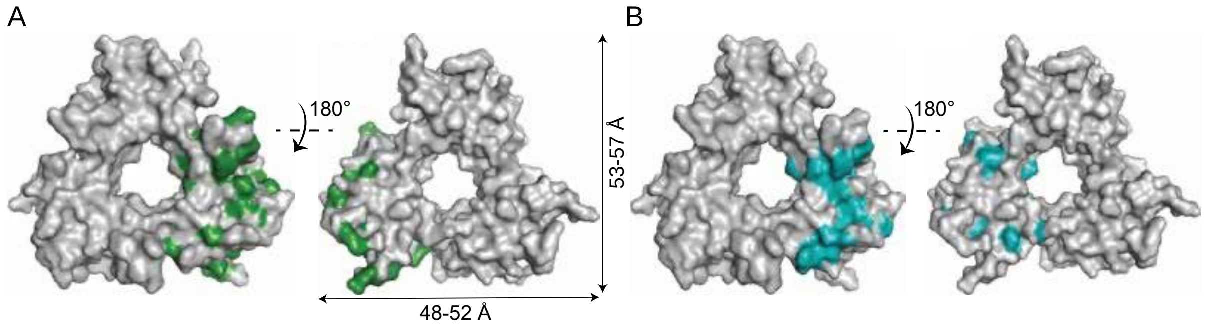


**Figure S7. Surface structure of SsgB.** (A) Key SsgB mutations are mapped onto the SsgB trimer structure (labeled green). Dimensions indicated by arrows are based on the protein backbone. (B) Evolutionary Conserved residues are highlighted in cyan on the SsgB trimer structure.

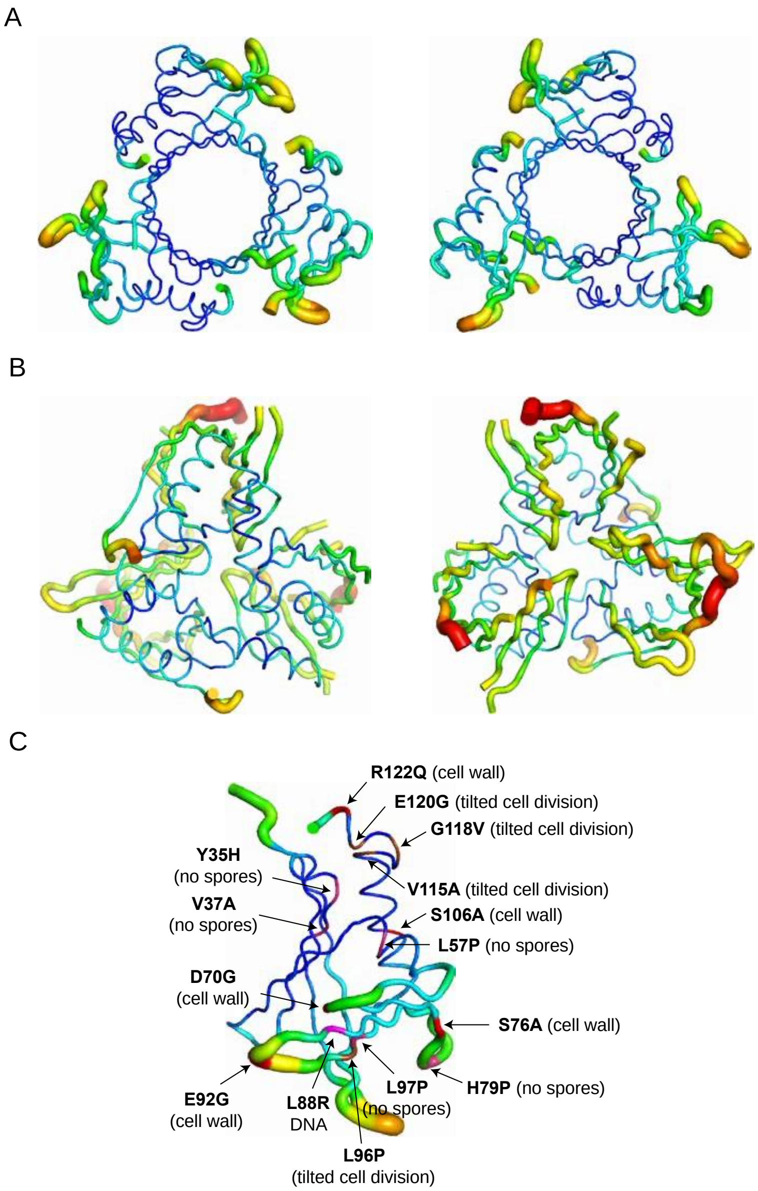


**Figure S8. Comparison of the B-factors for *Sc*SsgB and *Tf*SsgB and location of key mutations.** (A) (B) *Left,* top view of the B-factor putty structures of *Sc*SsgB and *Tf*SsgB (PDB code: 3CM1), respectively. *Right,* bottom view of the *Sc*SsgB and *Tf*SsgB B-factor putty structures. (C) Mutants labeled in the *Sc*SsgB B-factor putty structure. Most substitution mutants correlate to the formation of ectopic rotated septa (E120G, V115A, G118V, L96P), are localized in the low B-factor region, except residue L96P.

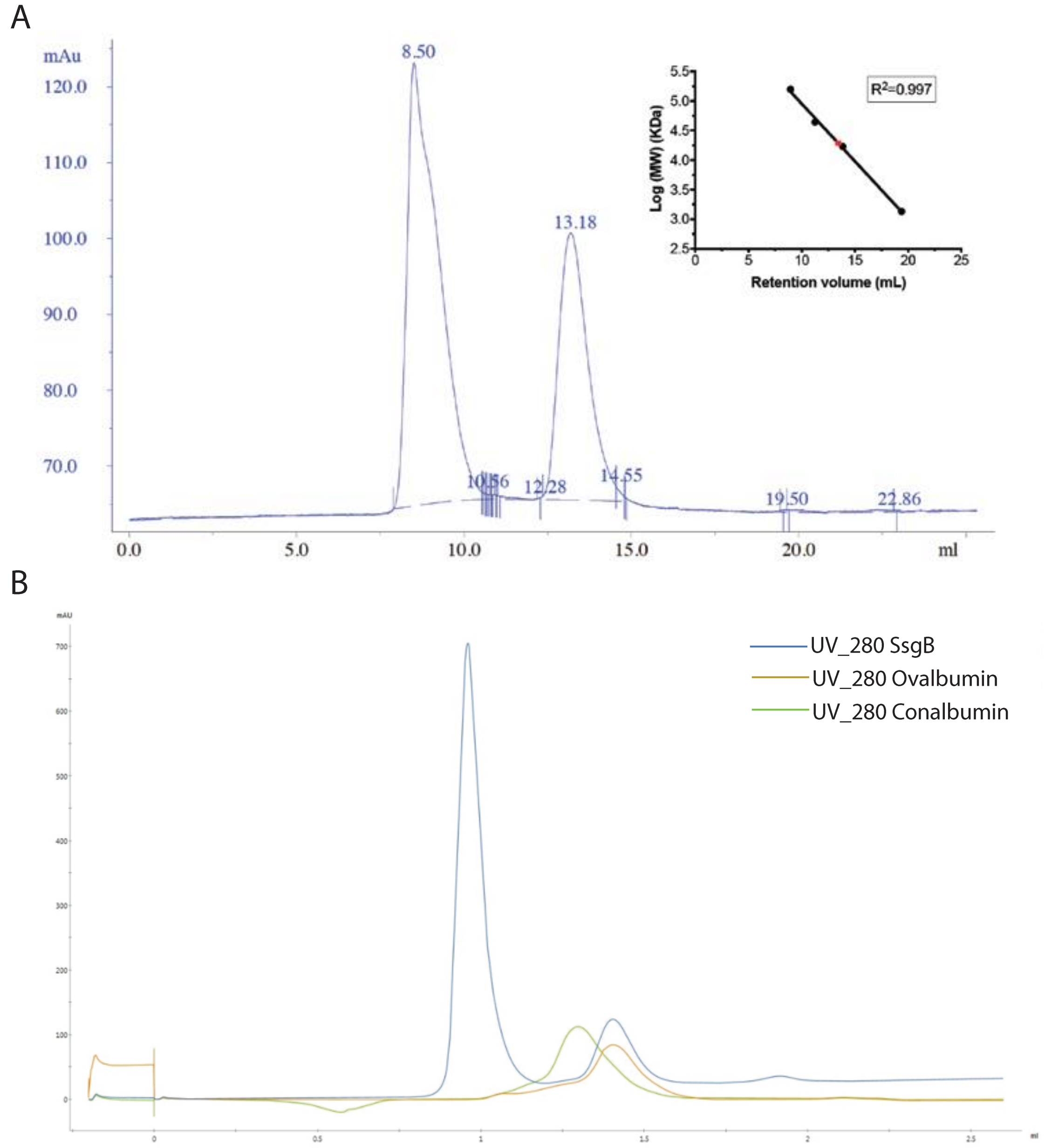


**Figure S9. Oligomerization studies of SsgB.** (A) Size-exclusion chromatography (SEC) analysis of SsgB (labeled as a filled red circle) shows primarily monomeric SsgB in solution. (B) On a 2.4ml Superdex 75 gel filtration column SsgB elutes at the same retention volume as Ovalbumin (44 kDa), consistent with SsgB trimers. A large peak at the void volume of the column (0.9 ml) indicates aggregated protein (MW: SsgB 15 kDa, Ovalbumin 44 kDa, Conalbumin 75 kDa).

**Table S1. Overview of sporulation phenotypes resulting from single amino acid substitutions in SsgB.** Residues that are important for SsgA function[1] are marked with an asterisk, those that are conserved between SsgB from different actinomycetes[2] are underlined.

| No sporulation (white) | Less sporulation (light grey) | Abundant sporulation (grey) |
|---|---|---|
| Y35H#, V37A#, L57P#, H79P, L97P#, E136G | S16P, T31A, L62P, D70G, S76A, V83A, E92G, S106A, V115G, Q128R, S131A, *138W | V15A, E18G, D30Y, T31M, H38R, V49G, W51*#, W51R#, D56G, H63L, T66A, E94G, L96R#, L96P#, E105G, G118V, E120G |

**Table S2. Overview of spore sizes of single SsgB mutations.** Those in grey fields are the wild type amino acid, W is white mutant, boxes show enlarged spores and black represent small spores. Solid white squares mark mutations that were not obtained in the library.

| | Amino acid | | | | | | | | | | | | | | | | | | | | | |
|---|---|---|---|---|---|---|---|---|---|---|---|---|---|---|---|---|---|---|---|---|---|---|
| Position | A | C | D | E | F | G | H | I | K | L | M | N | P | Q | R | S | T | V | W | Y | # of mutations | # of Whites |
| 51 | 1.40 | 1.37 | 1.46 | | W | 1.62 | W | W | W | W | W | 1.67 | 1.30 | | W | W | W | 1.35 | 1.54 | 1.42 | 21 | 9 |
| 88 | | | | | 1.45 | 1.34 | 1.47 | W | | 1.48 | 1.54 | W | | | 1.75 | 1.36 | 1.53 | 1.62 | W | 1.55 | 16 | 3 |
| 95 | 1.47 | 1.58 | 1.38 | W | W | 1.40 | | 1.54 | W | 1.36 | W | 1.57 | W | W | W | W | W | W | W | 1.37 | 22 | 11 |
| 96 | | 1.38 | 1.40 | 1.41 | 1.50 | 1.58 | 1.50 | 1.50 | 1.42 | 1.53 | 1.39 | 1.53 | | 1.37 | 1.44 | 1.65 | 1.50 | W | 1.49 | 1.36 | 21 | 1 |
| 97 | 1.49 | 1.54 | W | W | 1.42 | 1.48 | W | | W | 1.39 | | W | W | W | 1.43 | 1.44 | 1.51 | W | | 1.40 | 20 | 8 |
| 115 | 1.74 | | 1.34 | W | W | 1.30 | W | 1.65 | 1.53 | 1.60 | 1.35 | 1.34 | W | W | 1.41 | W | 1.49 | 1.49 | 1.48 | 1.45 | 22 | 6 |
| 116 | 1.49 | 1.41 | 1.31 | 1.67 | | 1.42 | 1.34 | W | 1.47 | 1.49 | | 1.36 | 1.67 | 1.47 | W | W | 1.49 | 1.66 | 1.40 | 1.51 | 21 | 3 |
| 117 | 1.44 | | | W | W | | 1.40 | 1.43 | W | W | 1.41 | 1.48 | 1.48 | 1.77 | W | 1.70 | 1.68 | W | W | 1.33 | 20 | 7 |
| 118 | 1.56 | | | | 1.42 | 1.33 | W | 1.40 | 1.50 | 1.31 | 1.43 | 1.35 | 1.51 | 1.55 | 1.28 | 1.37 | 1.64 | 1.51 | 1.43 | | 19 | 1 |
| 119 | | | | | | 1.45 | | | 1.41 | 1.43 | | | | | 1.49 | 1.36 | 1.49 | | 1.35 | | 10 | 0 |
| 120 | 1.59 | W | | 1.43 | 1.38 | 1.69 | 1.34 | 1.53 | 1.55 | 1.45 | | 1.43 | 1.79 | 1.47 | 1.45 | 1.48 | 1.42 | | | | 18 | 1 |
| 121 | 1.53 | 1.70 | | 1.51 | | | 1.52 | 1.75 | 1.35 | 1.61 | 1.30 | | | 1.38 | 1.51 | | 1.40 | 1.52 | 1.34 | | 16 | 0 |
| 122 | | 1.55 | 1.43 | | 1.56 | 1.45 | 1.45 | 1.51 | 1.47 | 1.48 | 1.53 | 1.50 | 1.54 | 1.86 | 1.38 | 1.59 | 1.38 | | | 1.33 | 19 | 0 |
| 123 | 1.58 | | 1.44 | 1.55 | 1.65 | 1.49 | 1.39 | | 1.50 | 1.46 | 1.41 | 1.40 | 1.58 | 1.49 | 1.54 | W | 1.42 | 1.38 | 1.44 | 1.38 | 21 | 1 |
| 124 | 1.39 | | | 1.48 | 1.36 | 1.35 | 1.50 | 1.41 | | 1.32 | W | | 1.66 | 1.34 | 1.30 | 1.39 | 1.37 | 1.54 | 1.34 | 1.52 | 19 | 1 |
| 125 | | | 1.53 | 1.49 | W | 1.31 | | | 1.51 | 1.41 | 1.41 | 1.51 | | | | 1.46 | 1.54 | W | 1.79 | | 15 | 2 |
| 126 | | | | | | | 1.52 | | | 1.62 | 1.61 | 1.40 | W | 1.32 | | 1.71 | 1.44 | 1.48 | 1.38 | 1.36 | 14 | 1 |
| 127 | 1.37 | 1.39 | 1.41 | 1.62 | 1.36 | 1.40 | 1.48 | 1.61 | 1.28 | 1.48 | 1.47 | 1.35 | 1.31 | 1.33 | 1.42 | 1.60 | | 1.47 | 1.28 | 1.40 | 22 | 0 |
| 128 | 1.47 | 1.57 | | 1.73 | 1.41 | 1.47 | 1.48 | | 1.50 | 1.47 | 1.47 | | W | 1.37 | 1.50 | 1.71 | 1.53 | 1.38 | W | 1.45 | 20 | 2 |
| 129 | 1.43 | 1.56 | 1.30 | 1.45 | | | | 1.60 | 1.52 | | | | | | | 1.58 | 1.59 | | | | 11 | 0 |
| 130 | | | | | 1.51 | 1.44 | 1.53 | 1.65 | | 1.41 | 1.34 | | | 1.45 | | 1.46 | 1.52 | 1.72 | 1.39 | | 14 | 0 |
| 131 | W | 1.42 | | | | 1.39 | | 1.46 | 1.67 | 1.54 | 1.67 | 1.53 | 1.33 | 1.89 | 1.34 | 1.50 | 1.72 | | 1.36 | 1.55 | 18 | 1 |

**Table S3. Summary of rotated septation observed in SsgB E120 substitution mutants.**

| Mutant | Normal septa | Rotated septa | Longitudinal septation | Ratios (%) | |
|---|---|---|---|---|---|
| | | | | Rotated | Longitudinal |
| **E120A** | 1123 | 39 | 6 | 3.5 | 0.5 |
| **E120C** | 1257 | 60 | 6 | 4.8 | 0.5 |
| **E120D** | - | - | - | - | - |
| **E120E** | - | - | - | - | - |
| **E120F** | 258 | 20 | 8 | 7.8 | 0.3 |
| **E120G** | 1204 | 72 | 6 | 6.0 | 0.5 |
| **E120H** | 631 | 37 | 10 | 5.9 | 1.6 |
| **E120I** | 362 | 31 | 1 | 8.6 | 0.3 |
| **E120K** | 987 | 78 | 9 | 8.0 | 0.9 |
| **E120L** | 875 | 35 | 8 | 4.0 | 0.9 |
| **E120M** | - | - | - | - | - |
| **E120N** | 864 | 30 | 3 | 3.5 | 0.4 |
| **E120P** | 771 | 31 | 4 | 4.0 | 0.5 |
| **E120Q** | 892 | 37 | 10 | 4.1 | 1.1 |
| **E120R** | 58 | 2 | 0 | 3.4 | 0 |
| **E120S** | 449 | 20 | 4 | 4.5 | 0.9 |
| **E120T** | 526 | 37 | 11 | 7.0 | 2.1 |
| **E120V** | - | - | - | - | - |
| **E120W** | - | - | - | - | - |
| **E120Y** | - | - | - | - | - |
| **WT** | 1011 | 0 | 0 | 0 | 0 |
| **Totals** | 10257 | 529 | 86 | 5.2 | 0.8 |

**Table S4. Overview of site-saturated mutagenesis of key SsgB residues.**

| Site-saturated mutagenesis | Phenotype | Ratios (%) |
|---|---|---|
| Residues 51, 88, 95-97 | Non-sporulating | 38 |
| Substitution of L96 | Disturbed DNA condensation | 66.7 |
| Substitution of L96 by I, R, S | Rotated septation | 1.7 |
| Substitution of E120 by A, C, F, G, H, I, K, L, N, P, Q, R, S, T | Rotated septation | 5.2 |
| Substitution of E120 by A, C, F, G, H, I, K, L, N, P, Q, S, T | Longitudinal division | 0.8 |

**Table S5. Site-saturated mutagenesis of key SsgB residues and their effects on cell morphology and structures.**

| Key SsgB mutagenesis | Localization | Effects on morphology | Effects on the structure |
|---|---|---|---|
| **E120G** | α2- α3 region | septum rotation | Disrupts the salt bridge with R55, hydrogen bond with E120, T119, G118 and V115; breaks the π-π interaction with Y35, thus, affects the stability of α3. |
| **G118V** | α2- α3 region | septum rotation | Disrupts the hydrogen bond with E120 and P116, and thereby affects the stability of α3. |
| **V115G** | α2- α3 region | septum rotation | Disrupts the hydrogen bond with H121, the salt bridge with R55 and thereby affects the stability of α3. |
| **L96P** | Solvent exposed β6-β7 region | septum rotation | Breaks β7 and affects the stability of α2, which further disturbs the R55-E120 interaction. |

**Tabel S6. Strains used in this study and their characteristics.**

| Strain | Characteristics | Reference |
|---|---|---|
| *E. coli* JM109 | see reference | [3] |
| *E. coli Rosetta™ 2 (DE3) pLysS* | see reference | [4] |
| *E. coli* ET12567 | *dam, dcm, hsdM, hsdS, hsdR, cat, tet* | [5] |
| M145 | derivative of *S. coelicolor* A3(2) lacking plasmids SCP1 and SCP2 | |
| GSB1 | M145Δ*ssgB* | [7] |
| GSB1+B | GSB1 harbouring a construct based on pHJL401 that expresses wild-type SsgB from the native *ssgB* promoter | This study |
| GSB1+B(N##N) | GSB1 harbouring a construct based on pHJL401 that expresses SsgB variants from the native *ssgB* promoter | This study |
| GSB1+B(G118V-eGFP) | GSB1 harbouring a construct based on pHJL401 that expresses SsgB-eGFP (G118V) from the native *ssgB* promoter | This study |

**Tabel S7. Primers used in this study.**

| **Primers** | **Sequence (5' -> 3')** |
|---|---|
| SsgB_rv | CTAGAAGCTTTGTGTGCCGTATGCGGTTGTCC |
| SsgB_fw | CTGAGAATTCATATGAACACCACGGTCAGCTGC |
| pSsgB_fw | AAACGCCGACCTGCCAGCTCAG |
| pSsgB_rv | CTAGAAGCTTGGATCCCCGTGGTGTTCATATGCGCCAGG |
| GFP_rv | gactctagattacttgtacagctcgtccatgccg |
| SsgB_E120A_rv | gtcaAAGCTTaGCTTTCCGCCAGGATGTGCGAGAGCTCCTGATCGAGATCGAAGTGCCGGTGCGCCGTGCCGGGGGGCACGGCGGCGTCTGTGCG |
| SsgB_ΔC_rv | gtcaAAGCTTttaGGCGGCGTCTGTGCGCTTCAGGAAGG |
| FtsZ_fw | ctgtgcaaCCATGGTTGCAGCACCGCAGAATTAT |
| FtsZ_rv | ctgtgcaaCTCGAGTTTCAGAAAATCCGGAACGT |
